# Early Life Stress induces brain-wide electrical network predisposition to migraine

**DOI:** 10.1101/2024.11.14.622885

**Authors:** Micah Johnson, Maureen Eberle, Ian Hultman, Xinyu Zhang, Yassine Filali, Benjamin Hing, Molly Matkovich, Alli Jimenez, Sara Mitchell, Anjayooluwa Adegboyo, Radha Velamuri, Julia Miller, Kung-Sik Chan, Sanvesh Srivastava, Rainbo Hultman

## Abstract

**Background:** Migraine is a disorder of severe, recurrent headaches and debilitating sensory, cognitive and affective symptoms, often triggered by stress. Early life stress in childhood has been shown to increase the likelihood of migraine in adulthood in humans. Calcitonin-gene relate peptide (CGRP) has been shown to reliably and acutely induce migraine or migraine-like behavior in both humans and rodent models. Here we investigate the impact of early life stress and CGRP on migraine-related neural circuitry, as well as the impact of early life stress on CGRP-mediated migraine-like activity in order to better understand the mechanisms by which early life stress predisposes neural circuitry to migraine brain activity.

**Methods:** We implemented an early life stress paradigm in the outbred strain of mice, CD1. We evaluated the impact of peripheral CGRP on migraine-like behavior and employed multi-site in vivo neurophysiology in freely behaving mice. A changepoint analysis was used to dissect differences in individual CGRP-induced responses.

**Results:** We found that early life stress exacerbated migraine-related behavioral and network physiology. CGRP alone caused disruptions in neural oscillatory activity across a network of brain regions including the anterior cingulate cortex (ACC), amygdala (AMY), thalamus (Po, VPM, and MDthal), and parabrachial nucleus (PBN). We found that power across the network was lowered within 10 minutes of peripheral CGRP exposure, which was sustained for ∼40-50 min. Coherence was mostly disrupted in amygdalar brain region pairings, and took on a shorter timecourse, with partial rescue of these responses by migraine abortive, sumatriptan. We found that early life stress exacerbated most of these responses, especially AMY-thalamic coherence pairings, although early life stress in the absence of CGRP demonstrated no impact on the network overall. We further identified individual mice with brain-network activity hypersusceptible to migraine.

**Conclusions:** Our findings demonstrate that early life stress confers vulnerability to migraine, simultaneously impacting behavior and brain network activity responses to peripheral CGRP.

## BACKGROUND

Migraine is an intricate and incapacitating neurological disorder and the second leading cause of global disability ^1^. Migraine is characterized by recurrent headache episodes spanning 4 to 72 hours with various sensory anomalies including hypersensitivities such as photophobia, phonophobia, osmophobia, and allodynia ^2^. It is important to recognize that migraine is far more involved than just headache, and frequently occurs over characteristic stages including prodrome, aura (impacts ∼20% of individuals with migraine), headache, postdrome, and interictal phases ^3^. In addition to sensory hypersensitivities, many of these phases of migraine can include changes in perception, cognition, autonomic nervous system regulation, and mood, pointing toward multiple sites of regulation and impact, especially across the brain. While the underlying origin of migraine has been historically debated in terms of central and peripheral mechanisms, there is growing consensus of both peripheral and central contributors to migraine^4^. Regardless of origin, it is clear that migraine engages multiple functional domains (cognition, perception, emotion/affect etc.) that are known to have causal origins in the brain.

Beyond the complex symptomology of migraine, environmental factors, especially those that have been experienced early in life, can shape susceptibility to migraine in adulthood. Early life stress (ELS), which is defined by adverse events experienced during childhood, is associated with increased risk for chronic pain conditions, including migraine ^5^. Several epidemiological studies have indicated that experiencing ELS leads to a greater risk of migraine and headache severity during adulthood ^6-8^. ELS is also one of many possible mechanisms linking the comorbidity of migraine and mood disorders, such as major depressive disorder ^9^. Finally, ELS has been associated with altered connectivity in brain regions associated with emotional regulation and stress, which are also implicated in migraine (especially amygdala)^10,11^. However, the mechanisms by which ELS predisposes one to migraine remains unknown.

Clues to brain mechanisms engaged in migraine stem largely from a combination of symptomology and imaging studies. Imaging studies have revealed altered functional connectivity across brain networks in migraine patients as well as structural differences in pain-related regions of migraine patients (Figure 8A). These changes have been observed not only during migraine attacks but also in the interictal period ^3,12-14^. A number of studies have indicated that a large part of what makes migraine so severe and debilitating comes from the aversive aspect of pain, which is greater for pain in the head compared with pain in the periphery ^15-17^. Head pain is ranked as more emotionally salient and severe than other types of peripheral pains. For example, a study in humans demonstrated that repeated application of noxious heat to the face induces sensitization whereas the same application of noxious heat to the hand induces habituation ^15^.

Thus, we focused our initial efforts toward understanding endogenous neural activity of migraine across several brain regions with an emphasis on regions involved in the emotional salience of head pain and mood aspect of migraine and involvement in ELS: anterior cingulate cortex (ACC), amygdala (AMY), posterior thalamus (Po), ventroposteromedial thalamus (VPM), medial dorsal thalamus (MDthal), and parabrachial nucleus (PBN), all of which have been implicated in the pathophysiology of migraine ^3,18^. The ACC is a key region in pain processing and more specifically, emotional processing of pain ^19^. Reduction in ACC grey matter is correlated with frequency of migraine attacks in migraine patients ^20^. AMY is implicated in affective pain processing and the perception of such pain ^21,22^. Resting functional connectivity patterns in a network involving the amygdala have been shown to discern migraine and healthy control patients ^23^. Further, increased connectivity patterns between the nucleus accumbens and AMY were found during the preictal phase compared to the ictal phase in specific phases of migraine^24^. The thalamus is broadly implicated in the integration of sensory information and has been associated with migraine-related allodynia, photophobia, and hypersensitivity to negative emotional stimuli ^25-28^. Additionally, altered thalamo-cortical functional connectivity patterns have been reported in migraine ^29^. Optogenetic stimulation of the posterior thalamus was found to induce photophobia-like behavior in the absence of anxiety ^30^. Finally, the brainstem as a whole, and the PBN in particular, appear to play an important role in migraine^31,32^. PBN is a critical sensory relay center and is important for homeostatic function, arousal, cravings, thirst, nociception and regulation of affective responses ^33,34^. Altered connectivity patterns across subcortical-cortical networks that include the PBN have been found in migraine patients during the preictal phase ^12,18,35,36^. Importantly, the PBN contains CGRP projections to the central amygdala, which has been demonstrated to play an important role in the emotional aspect of pain ^21,33,37^ as well as projections to the thalamus, which convey nociceptive information to higher order cortical regions ^38^. Overall, human imaging studies point to brain-wide connectivity changes in migraine, however, ultimately endogenous multi-region brain network studies in disease models are required to understand the ways in which brain regions communicate on a timescale of neuronal activity.

The ways in which endogenous brain-network activity is engaged and coordinated in mouse models of migraine remains largely unknown. Preclinical models of migraine allow for probing of neuronal circuitry across time in a way that can help better understand the actual mechanisms underlying the interconnected circuitry that leads to migraine and to directly test the impact of prior exposure of ELS on the development of migraine-like brain activity and behavior. Developing an understanding of systems-level brain activity in preclinical models of migraine holds great promise for a new era of migraine research and therapeutic development ^39^.

One of the best characterized and most robust animal models of migraine makes use of calcitonin gene-related peptide (CGRP) to induce a migraine-like phenotype in mice. This model is especially important given CGRP therapeutics have played such an important role in migraine treatment and prevention, being the first pharmacologics designed specifically for migraine^40^. The CGRP model demonstrates validity to the human disorder in three important ways: face validity (the model displays a similar disease phenotype as in humans) ^41^ construct validity (the model shares the same underlying cause of the disease in humans) ^42,43^, and predictive validity (treatments that are effective in humans are effective in the model) ^42,44^. CGRP has emerged as a neuropeptide of prominent importance in migraine pathophysiology. Key evidence for the involvement of CGRP in migraine is that CGRP levels are elevated during migraine attacks ^45-47^. Blocking CGRP signaling can also alleviate migraine symptoms in both humans and rodent models ^48^. Additionally, intravenous infusion of CGRP (either central or peripheral) leads to a delayed migraine-like headache in humans ^43^ and preclinical migraine-like phenotypes (e.g., light aversion, spontaneous pain, and allodynia) in rodents ^41^. There has been mounting evidence that CGRP acts on receptors peripherally, with indirect impacts that reach the brain ^4,39^. CGRP has also been shown in preclinical models to evoke a migraine-like response when injected directly into the brain ^30^. However, the nature of the impact of peripheral CGRP on the brain remains one of the fundamental questions in solving the mystery of migraine.

In this manuscript, we aim to investigate the impact of ELS on CGRP-induced migraine-like activity in mice. To better understand brain network dynamics underlying migraine, we employ multi-site *in vivo* neurophysiological recordings in freely behaving animals to monitor changes in brain activity and connectivity in response to peripheral CGRP administration over time, providing both high spatial and temporal resolution. We then further probe brain network activity with anti-migraine medication, sumatriptan, and ELS, in order to understand how ELS may predispose brain networks to migraine. Finally, we applied a changepoint analysis in order to identify network-wide changes associated with migraine-like phenotypes.

## METHODS

### EXPERIMENTAL MODEL AND STUDY PARTICIPANT DETAILS

#### Animal Care and Use

All experiments were performed with approved protocols by the Institutional Animal Care and Use Committee (IACUC) of the University of Iowa. CD1 mice purchased from Charles River Laboratory were utilized for all experiments. For neurophysiological experiments (excluding maternal separation stress), male and female mice aged 9 weeks of age were group housed 2-3 animals per cage for 1 week prior to surgery. Mice were maintained on a 12 h light/dark cycle with ad libitum access to food and water. All testing was performed between 9:00 AM and 9:00 PM.

#### Maternal Separation Stress with Early Weaning and Limited Nesting

CD1 mice were obtained from Charles River Laboratory and bred in-house to generate breeding pairs and their offspring. All experimental animals were a minimum of one generation removed from shipping stress. Behavioral testing was conducted across several cohorts based on parturition date. The maternal separation stress paradigm was adapted from previously described methods ^49,50^. All mice were housed on beta chip bedding with only 1/3 of the supplemental nesting material provided to controls (Enviro-Dri nesting material). Litters were culled to 8 pups on P1 with balanced sex distribution. Starting at P2, pups were separated from dams for 4 hours daily until P5 then 8 hours daily until P16. Supplemental heat was provided during separation periods. Pups were weaned early at P17 to eliminate the likelihood of overcompensation of maternal care behavior upon dams reunion with pups ^51^. Prior to experiments, animals were weighed weekly and placed into balanced treatment groups based on weights and cohort.

### Neurophysiological Experiments

#### Drug Administration

All injections were performed intraperitoneally (IP) with a 25G needle. We administered calcitonin gene-related peptide (CGRP) (Sigma-Aldrich, Saint Louis, MO) at 0.1 mg/kg and sumatriptan at 0.6 mg/kg (AuroMedics Pharma LLC, E. Windsor, NJ). We utilized phosphate-buffered saline (PBS) as a diluent and vehicle for all experiments. We co-injected sumatriptan and CGRP by doubling the total volumes of each and diluting them such that our co-injections were the same solution volume as the single CGRP and PBS injections. All solutions were prepared 1 hour prior to the start of experiments and placed on ice for the remainder of the experiment.

#### Electrode Implantation

Animals aged 10-12 weeks were anesthetized with isoflurane, placed in a stereotaxic device, and craniotomized for electrode implantation. Metal ground screws were secured above the olfactory bulb and cerebellum. Tungsten wires (California Fine Wires) were implanted in the anterior cingulate cortex (ACC), ventral posteromedial thalamus (VPM), medial dorsal thalamus (MDthal), posterior thalamus (Po), central amygdala (AMY), and left and right parabrachial nucleus (LPBN/RPBN). Coordinates were adapted for CD1 animals and verified histologically. Male surgical coordinates for the center of bundles placed (not actual electrode tips) are as follows: ACC, 0.8mm A/P, 0.25mm M/L, 1.8mm D/V; Amy (BLA/CeA) -1.58mm A/P, 2.75mm M/L, 3.9mm D/V; thal (MDthal/Po/VPM) -1.82mm A/P, 1.5mm M/L, 3.1mm D/V; PBN - 5.5mm A/P, +/-1.5mm M/L, 2.3mm D/V. Female are: 0.78mm A/P, 0.24mm M/L, 1.8mm D/V; Amy (BLA/CeA) -1.54mm A/P, 2.68mm M/L, 3.8mm D/V, thal (MDthal/Po/VPM) -1.77mm A/P, 1.46mm M/L, 3.02mm D/V; PBN -5.36mm A/P, +/-1.46mm M/L, 2.24mm D/V.

#### Light-Aversion Test

To assess light-aversive behavior, we performed a light/dark assay. In this assay, animals were placed in a transparent testing chamber (MedAssociates, East Fairfield, VT) consisting of a dark or light zone divided by a dark insert. We utilized infrared beam tracking and Activity Monitor software to analyze activity across a 30-minute time period under 20,000 lux light intensity. While a range of time windows for the light/dark assay have been used across the literature, in our hands, 15 minutes has always provided the best metric of CGRP response, even when compared against a 30-minute test (Figure S2), which is consistent with other studies in the literature ^42,52^. Because some studies also report CGRP-induced behavioral changes across 30 min, we also collected additional data beyond the 15-minute assessment window in order to be thorough. On the first day, animals underwent a baseline assessment in which they are assessed for a 15-minute period. Mice were injected with vehicle the following day and assessed for another 15-minute period. Lastly, animals were injected with CGRP and activity was assessed for 15 minutes in the testing chamber. For ELS experiments, testing occurred at P81-85.

#### Neurophysiological recordings

Mice were connected to Mu headstages (Blackrock Microsystems, UT, USA) under isoflurane anesthesia prior to recordings. During recordings, mice were placed in a homecage environment consisting of standard cages with corn cob bedding and recorded for 30 minutes under baseline conditions. Neurophysiological data was captured using the Cereplex Direct acquisition system and sampled at 30kHz. Local field potentials (LFPs) were stored at 1000 Hz and a bandpass filter was applied at 0.5-250 Hz.

#### Timeline of neurophysiological recordings during peripheral administration of CGRP

Two weeks following recovery from surgery, animals underwent three days of homecage recordings. On Day 1, animals were recorded for 30 minutes then received an IP PBS vehicle injection and recorded for an additional 2 hours post-injection. Day 2, we administered IP CGRP injections. To ensure any observed changes in brain activity on Day 2 could be attributed to changes induced by CGRP and not the injection itself, we repeated an IP PBS vehicle injection recording on Day 3.

#### Histology/Electrode Placement

All animals were euthanized upon completion of experiments and perfused with 4% paraformaldehyde (PFA). Brains were stored in 30% sucrose overnight and submerged in an OCT compound then frozen on dry ice and stored at -80°C. Brains were sliced in 35µm sections then washed in PBS prior to being stained with red Nissl (ThermoFisher, Waltham, MA). Slices were imaged at 5x magnification using a Leica fluorescence microscope. Each individual slice was aligned to the Paxinos and Franklin brain atlas for verification of electrode placement. Overall, electrode placements had a ∼97% success rate. Two animals were removed from analysis due to failure to hit targets.

### Early Life Stress Behavioral Experiments

#### Elevated Plus Maze

To assess anxiety-like behavior in mice, we utilized an elevated plus maze (EPM) apparatus consisting of two opposing open and closed arms. A video camera was placed directly above the center of the maze to capture activity during each trial. Each animal was placed in the center of the EPM facing the same closed arm at the start of each trial and tested for a total of 5 minutes. All mice were tested at P66-69.

#### Plantar von Frey Test

To assess mechanical allodynia in mice, we utilized the plantar von Frey test. Over a total of 5 days, mice were placed in acrylic chambers over a grid support (Bioseb, France). Mice habituated to the chambers for a total of 2 hours the first two days. For each of the following days, mice habituated to the chambers for 1 hour prior to the start of experiments. Mice were considered well habituated when standing on all four paws, calm, and non-responsive to the habituation filament. Starting on Day 3 (Baseline day), a von Frey filament was applied to the plantar surface of the right hindpaw. We followed the “up-down” method to obtain a paw withdrawal threshold for each animal ^53,54^. On Day 4, all animals received an IP PBS (vehicle) injection and were allowed to rest for 30 minutes prior to testing. On Day 5, animals received either CGRP or vehicle injections with the administering experimenter blinded to conditions, and were allowed to rest for 30 minutes prior to testing. Animals were tested at P72-79.

#### Open Field Test

To determine whether the light aversive behavior was independent of anxiety-like behavior in mice, we performed an open field test. Animals were tested at the end of the experiments at P83-86 the day after light aversion experiments. Mice were placed in a large open field box for 20 minutes at a time following administration of vehicle or CGRP, with the administering experimenter blinded to conditions. We utilized ANY-maze software to monitor the total amount of time animals spent in the center zone of the open field box.

Light aversion test for ELS was carried out as described above with the administering experimenter blinded to conditions.

### QUANTIFICATION AND STATISTICAL ANALYSIS

#### Implanted vs. Unimplanted Behavioral Statistics

We fit a three factor mixed effects model using the *lme4 R* library ^55^ with fixed effects terms for the model intercept, CGRP, day, implant and the interaction between CGRP and implant. We employed a random intercept for each mouse. We obtained the reported p-values for the fixed effects terms using the *lmerTest R* library ^56^.

#### Directionality Analysis

LFP signal directionality between pairs of brain regions was estimated using a temporal lag approach introducing phase offsets as previously described ^57^. Butterworth bandpass filters were applied to isolate LFP oscillations within 1Hz windows using a 1Hz step in the 1 50Hz range. The Hilbert transform was used to determine the instantaneous phase of the filtered LFPs and the instantaneous phase offset time series was calculated for each LFP brain region pair. The mean resultant length (MRL) for the phase offset time series was calculated, yielding the phase coherence between pairs of LFPs. Temporal offsets were introduced between each pair of LFPs ranging from 250ms to 250ms in 2 ms steps and the phase coherence was recalculated between each LFP pair.

#### Frequency Band Identification

In order to determine the specific frequency bands at which pairs of brain regions demonstrated clear temporal offset (leading or lagging), we used a fused lasso approach within the 1*Hz* to 50*Hz* range obtained from the phase offsets data for 34 mice and for each pair of brain regions from the set: ACC, LPBN, RPBN, CeA, BLA, MD_thal, Po, VPM. Lasso (<u>L</u>east <u>a</u>bsolute <u>s</u>hrinkage and <u>s</u>election <u>o</u>perator), is a model parameter selection and regularization method. The lasso method minimizes the least squares loss over a set of parameters in a linear regression problem by imposing the condition that the sum of the absolute values of the parameter estimates (that is, the 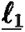 norm of the parameters) is less than some value ^58^. This method generally results in some parameter estimates being zeroed out, in effect selecting the most important covariates that are more predictive of the response than those with zeroed out regression coefficients. The fused lasso accounts for temporal or spatial dependence amongst covariates by further imposing a condition that the sum of the absolute values of the lag one differences between subsequent parameter estimates be less than some value ^59^. Modeling the response under fused lasso then

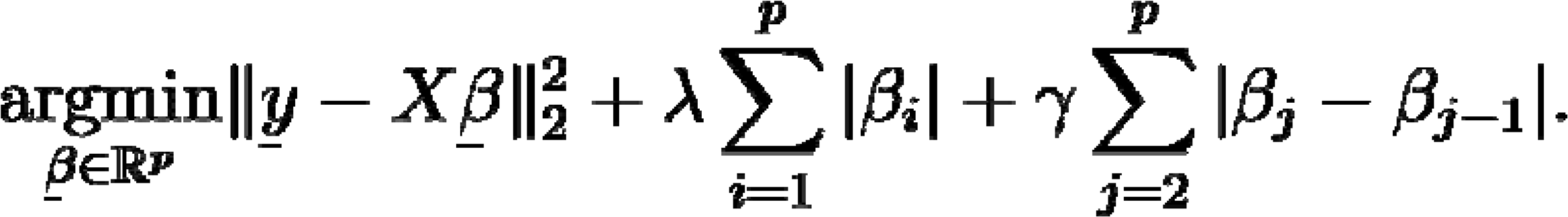

amounts to finding a set of parameter estimates 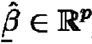 as a solution of The parameters 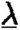 and 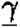 are positive tuning parameters. The solution of the optimization problem in (1) depends on the chosen value of 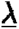 and 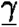, that is 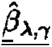, which are chosen via cross validation in such a way that the test mean squared error is minimized.

We hypothesized that a small number of frequencies are active in these measurements and an active frequency implies that neighboring frequencies are also active. We applied the application of fused lasso for signal approximation ^60^. We first computed the mean over all observations for each pair of brain regions and for each mouse. This resulted in 168 observations (8 mice times 21 brain region pairs) each with 50 elements corresponding to the mean phase offset at each frequency from 1*Hz* to 50*Hz*. This 168 × 50 matrix was then reshaped into a 400 × 21 response matrix **Y** in which each column contained the means across frequencies for each mouse for a particular pair of brain regions. The design matrix **X** was comprised of eight 50 × 50 identity matrices stacked on top of each other (one for each mouse) resulting in a 400 × 50 matrix. The 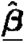 matrix of parameter estimates was a 50 × 21 matrix for which each column contained the model estimates for the phase offsets over the 1*Hz* to 50*Hz* range for a given pair of brain regions. We then used the genlasso library (*genlasso: Path Algorithm for Generalized Lasso Problems*, R package 1.5) in R to obtain estimates 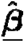 under the fused lasso model with the tuning parameters 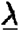 and 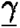 set to 3 and 0.5 respectively. This resulted in 21 different vectors (the columns of 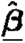) with bands of non-zero values corresponding to significant phase offset frequencies and zeros indicating non-significant frequencies for each pair of brain regions.

#### Frequency bands of interest

The Po-CeA relationship was the only brain region pairing that included a different, 1-7 Hz band. We accounted for 1-3 Hz and 1-7 Hz in our multiple comparisons corrections, however we did not observe substantial evidence for the biological relevance of the 1-3 Hz band across all brain regions we observed given how small the differences in this band were relative to the other frequency bands and the fact that it was only significant in one region pairing and not all (Supplemental Figure S3) so we focused on the 4-7 Hz and 8-13 Hz frequency bands.

#### Analysis of Power and Coherence over time

We applied a log transformation to the recordings to preprocess the power data. For every brain region and every mouse on each day of the experiment, we averaged the log-transformed power across all channels within each brain region and all frequencies within specific frequency bands. This resulted in a single log-power time series for every frequency band and for every brain region being associated with each mouse on each day of an experiment. We computed the mean log-power over baseline recordings for each frequency band and brain region for each mouse on each day of a given experiment. We normalized the post-injection log-power recordings relative to the baseline by subtracting these means from the corresponding log-power post-injection recordings.

We preprocessed the coherence data similarly. For each pair of brain regions and all frequencies within specific frequency bands, the coherence recordings were averaged across all the channel pairs, resulting in a single coherence time series for every frequency band across each mouse’s 28 pairs of brain regions on each day. We then applied a log-odds transformation to the averaged coherence data to normalize the post-injection log-odds coherence recordings using the same approach as we did for the power data. Specifically, we calculated the mean of the baseline log-odds coherence for each frequency band and brain region pair and subtracted these means from the corresponding post-injection log-odds coherence values.

After preprocessing, we used these data to fit the FOS model using generalized additive model (GAM) function in the mgcv library in R to plot 30 second rolling means in order to identify underlying trends over time ^61^. We first computed 30-second rolling means over each mouse’s log-power post-injection time series for each day of the experiment, for each frequency band and each brain region, using the *tf_smooth* function in the *R* library *tf* ^62^. Next, we averaged the vehicle 30-second rolling mean time series across days. This resulted in each mouse having one log-power 30 second rolling mean post-injection time series for each treatment (vehicle and CGRP), frequency band, and brain region. We used these series for all the mice to compute the average and one standard deviation and obtained one mean CGRP log-power time series, one mean vehicle log-power time series and their respective one standard deviation time series for each frequency band for each brain region. For each frequency band in each brain region, we plotted these 30-second rolling mean time series together to facilitate comparison between the CGRP and vehicle conditions. We used the *R* packages *ggplot2^63^*and *tidyfun^64^* to plot these figures.

We applied the same procedure to the coherence data, resulting in one mean CGRP log-odds coherence 30-second rolling mean time series, one mean vehicle log-odds coherence 30-second rolling mean time series, and their respective one standard deviation time series for each frequency band across each pair of brain regions. We plotted these 30 second rolling mean time series together for comparison in the same manner as the log-power analyses.

For the power GAM analysis, we computed the mean of each mouse’s log-power post-injection vehicle time series, resulting in one log-power post-injection time series for vehicle and one for CGRP for each frequency band in each brain region. We obtained one pairwise difference time series for each frequency band in each brain region for each mouse by subtracting the mouse’s log-power vehicle time series from the corresponding CGRP time series. For each frequency band in each brain region, we used 90 minutes of the post-injection recordings for the power data analysis.

The coherence analysis is performed similarly, except we computed the mean log-odds coherence instead of the mean log-power. This resulted in each mouse having one log-odds coherence post-injection time series for the vehicle and one for CGRP for each frequency band across each pair of brain regions. We then subtracted each mouse’s log-odds coherence vehicle time series from the corresponding CGRP time series to obtain one pairwise difference time series for each frequency band across each pair of brain regions for each mouse. For each frequency band in each brain region, we used 60 minutes of the post-injection recordings data for the coherence data analysis.

The coherence analysis is performed similarly, except we computed the mean log-odds coherence instead of the mean log-power. This resulted in each mouse having one log-odds coherence post-injection time series for the vehicle and one for CGRP for each frequency band across each pair of brain regions. We then subtracted each mouse’s log-odds coherence vehicle time series from the corresponding CGRP time series to obtain one pairwise difference time series for each frequency band across each pair of brain regions for each mouse. For each frequency band in each brain region, we used 60 minutes of the post-injection recordings data for the coherence data analysis.

For the power and coherence analyses, we used the *gam* function in the *R* package *mgcv* (Wood, 2011) for FOS regression. Each model had the following form:

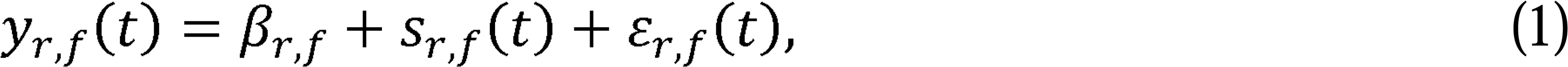

where *y*_*r,f*_(*t*) denotes the pairwise difference in log-power or log-odds coherence at time point *t,* brain region (for power) or pair of regions (for coherence) *r* and frequency band *f*; *β*_*r,f*_ denotes the (*r, f)*-specific model intercept; *s*_*r,f*_(*t*) denotes the *(r, f)*-specific model smooth function-over-time effect at time point *t*; and *ε*_*r,f*_(t) denotes *(r, f)*-specific independent centered Gaussian noise at time point t. This FOS regression model assumes that the pairwise differences in log-power or log-odds coherence are frequency band and region- or pair-specific functions of time.

We fit this model in *R* as follows. For the mouse *i*, *X*_*i,r,f*_ denotes the *R* data.frame with columns representing time and response, where *t* is the vector of evenly spaced time steps and *y* denotes the observed pairwise difference in log-power or log-odds coherence for mouse *i* in brain region or pair of brain regions *r* and frequency band *f*. Let *X_rf* denote the data.frame formed by stacking the elements of *X*_*i,r,f*_ on top of each other for *i* = 1, …., 34. We fit the FOS regression model using the *mgcv* package in *R* as follows:

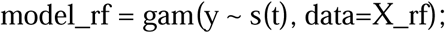

The fit model_rf contains the fitted values indicating the mean effect of CGRP over time along with p-values for both the intercept and function-over-time components *β*_*r,f*_ and *s*_*r,f*_(*t*) respectively. We plotted each model’s fitted values along with a two-standard error region. When these two-standard error confidence regions contain zero, we interpret this as indicating no significant CGRP effect at that point in time.

#### Analysis of Sumatriptan over time

For 10 of the 34 mice from the previous experiment, we collected additional power and coherence data over three more days. This n was determined for a repeated measures design in keeping with similar studies^65,66^. Each mouse received a vehicle injection on the first day, a Sumatriptan injection on the second day, and an injection of both CGRP and Sumatriptan on the third day. This resulted in each of these 10 mice having a total of three vehicle injection days, one CGRP-only injection day, one Sumatriptan-only injection day and one CGRP-with-Sumatriptan injection day. The recorded power and coherence data were preprocessed as described earlier, and each set of vehicle time series’ were averaged into one vehicle time series such that each mouse had a vehicle time series, a CGRP-only time series, a Sumatriptan-only time series and a CGRP-with-Sumatriptan time series associated with each frequency band and each brain region for power or pair of brain regions for coherence.

For both power and coherence, we again computed the 30-second rolling means for the post-injection time series using the *tf_smooth* function from the R library *tf* (Scheipl & Goldsmith, 2024)^62^ and plotted them in the same manner as in the previous experiment. Now, however, each plot has four lines: one for vehicle, one for CGRP, one for Sumatriptan and one for CGRP-with-Sumatriptan.

For the power and coherence FOS GAM analyses, we focused only on comparing CGRP to CGRP-with-Sumatriptan. These analyses follow the previous section. For every mouse, we obtained a pairwise power difference time series for each frequency band in each brain region by subtracting each mouse’s log-power CGRP-with-Sumatriptan time series from the corresponding CGRP-only time series. Similarly, for the coherence analysis, we obtained the pairwise difference time series for each frequency band across each pair of brain regions for each mouse by subtracting each mouse’s log-odds coherence CGRP-with-Sumatriptan time series from the corresponding CGRP-only time series. We used 45 minutes of the post-injection recordings data for our power analyses, and we used 20 minutes of the post-injection recordings data for our coherence analyses.

For the power and coherence analyses, we used a function-on-scalar regression using the *gam* function in the *R* package *mgcv* (Wood, 2011)^67^. Each model had the same form as equation (1) above and was fit in the same manner as the previous experiment. We plotted the fitted values along with two-standard error confidence bands and obtained each model’s p-values corresponding to the intercept and function-over-time components *β*_*r,f*_ and *s*_*r,f*_(*t*) respectively.

#### Analysis of Early Life Stress Data over Time

We collected power and coherence data from 11 different mice over three days. Similar to the experiment comparing CGRP to vehicle, each mouse received vehicle injections on the first and third days and a CGRP injection on the second day. Of these 11 mice, six were ELS mice and five were control mice. We preprocessed the power and coherence data as described above. This experiment differed from the previous two experiments in that each treatment group consisted of different mice.

We computed the 30 second rolling mean plots in a similar manner as the previous two experiments using the *tf_smooth* function in the *R* library *tf* (Scheipl & Goldsmith, 2024)^62^. For the power analysis we first computed 30 second rolling means over each mouse’s log-power post-injection time series. Next, we averaged each mouse’s vehicle 30-second rolling mean time series across days. This resulted in each mouse having one log-power 30 second rolling mean post-injection time series for each treatment (vehicle and CGRP), frequency band, and brain region. We grouped the observations by ELS vs. control and vehicle vs. CGRP and computed the average and one standard deviation log-power 30 second rolling mean time series across the observations within each group. This resulted in separate mean and one standard deviation log-power time series for each group (ELS-with-CGRP, ELS-with-vehicle, control-with-CGRP and control-with-vehicle) for each frequency band for each brain region. For each frequency band for each brain region, we plotted these 30 second rolling mean time series together for comparison in the same manner as the previous experiments.

We processed the coherence data in the same way so that we had separate mean log-odds coherence 30 second rolling mean time series for ELS-with-CGRP, ELS-with-vehicle, control-with-CGRP and control-with-vehicle along with their respective one standard deviation time series for each frequency band for each pair of brain regions. We plotted these 30 second rolling mean time series together for comparison in the same manner as the log-power analyses.

The different experimental design implied we use a different GAM model. We performed FOS GAM analyses for each frequency band in each brain region for power or pair of brain regions for coherence based on a two-factor model with interaction, where the first factor was ELS vs. control and the second factor was CGRP vs. vehicle. Because each mouse had observations over three different days, we used a mixed effects model with random intercept and smooth function-over-time terms for each mouse. Each model used either log-power or log-odds coherence as the response and had the following terms:

● Overall model intercept
● Mouse-specific random intercept
● Mouse-specific random smooth function-over-time
● Overall smooth function-over-time
● ELS effect intercept
● CGRP effect intercept
● ELS:CGRP interaction effect intercept
● ELS effect smooth function-over-time
● CGRP effect smooth function-over-time
● ELS:CGRP interaction effect smooth function-over-time

This FOS regression model assumes that the log-power or log-odds coherence are frequency band and region- or pair-specific functions of time. For both the power and coherence analyses we used 45 minutes of the post-injection recordings.

We fit this model in *R* as follows. For the mouse *i*, *X*_*i,r,f*_ denotes the *R* data.frame with columns representing time, ELS vs. control labels, CGRP vs. vehicle labels, the interaction between ELS and CGRP labels, the mouse ID and response. Let *t* denote the vector of evenly spaced time steps repeated for each day’s recordings, *els* denote the vector of factor labels indicating whether an observation is associated with an ELS or control mouse, *cgrp* denote the vector of factor labels indicating whether an observation is associated with CGRP or vehicle injection, *els_w_cgrp* denote the ELS:CGRP interaction dummy variable vector indicating whether an observation is both CGRP and ELS or not, *mouse* denote the mouse ID associated with the observation and *y* denote the observed log-power or log-odds coherence for mouse *i* in brain region or pair of brain regions *r* and frequency band *f*. Let *X_rf* denote the data.frame formed by stacking the elements of *X*_*i,r,f*_ on top of each other for *i* = 1, …., 11. We fit the FOS using the *mgcv* package in *R* as follows:

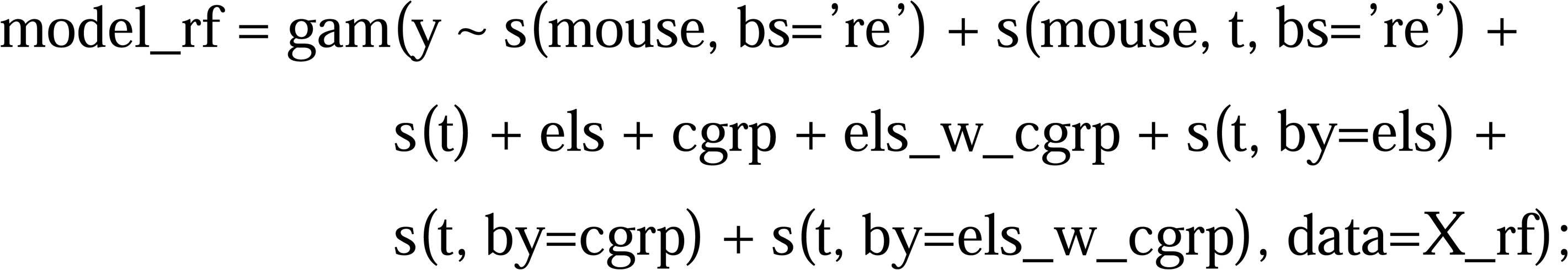

From the fit model_rf, we obtained the smooth function-over-time components of the fitted values which provide the estimated ELS effects over time and the estimated ELS:CGRP interaction effects over time. We plotted these estimated effects over time along with a two standard error confidence region. We also obtained from the model the p-values associated with the ELS effect intercept, the ELS effect smooth function-over-time, the ELS:CGRP interaction effect intercept and the ELS:CGRP interaction effect smooth function-over-time.

We took all of the p-values obtained from each experiment’s analyses and computed FDR corrections on them in order to perform multiple comparisons hypothesis tests of whether or not there was a significant difference in treatment effects over time for a given brain region or pair of brain regions in a particular frequency band.

#### Changepoint analysis

Changepoint analysis was carried out as originally described in Zhang & Chan (2024) ^68^. For this analysis, we evaluated the data in the two frequency bands we found to be most impacted by peripheral CGRP (4-7Hz and 8-13 Hz).

For conciseness, we shall simply describe the case of detecting changes in the 4-7 Hz component of the LFP, as the approach is the same for both frequency bands. Specifically, for each mouse, the 7-dimensional LFP times series are blocked into non-overlapping 10-second blocks, over each of which (i) we computed its smooth spectral density matrix estimate (as a function of frequency with 1 Hz steps) via a Bartlett window, (ii) normalized the spectral density matrix estimates so that their diagonal elements are identically equal to 1 and the off-diagonal elements measuring the LFP coherency between pairs of brain regions, and (iii) summarized the normalized spectral density matrix function by averaging over the 4-7 Hz.

The matrices per the 10-second blocks are 7 by 7 complex valued matrices, and their real part can be interpreted as proportional to the correlation matrix of the 7-dimensional 4-7 Hz (periodic) components of the LFP time series, with the latter denoted as Ξ_*t*_ for the t-th block (section 7.1 in Brillinger (2001)^69^. The change-point detection then proceeds to find the epochs (i.e. the change points, if they exist) across which Ξ_*t*_’s undergo a change, as follows. Let *v* denote the projection vector, of unit length, pointing the principal direction of the change in Ξ_*t*_, i.e., the quadratic form *v^*T*^*Ξ_*t*_ *v* changes the most across the change point, among any unit directional vectors. The projection v is then the principal direction along which the pre- and post-change-point correlation matrices of the 4-7 Hz periodic components of LFP differ from each other to the greatest extent.

The projection vectors reveal the structure of the change of the coherency matrix over the detected change point. Larger values in magnitude indicates larger changes for the corresponding brain regions. Opposite signs within the projection vector are indicative of two major types of organizational matrix changes, which can be identified as blocks within the change in coherency matrix across the changepoint. These blocks can segregate coherency into activity within a subgroup of brain regions as two groups and activity between the subgroups (as in Fig. 7E). If the projection comprises coefficients of opposite signs, the brain regions are split into two groups according to the signs and may signify that across the change point, the inter-brain-region correlations between the two groups change differently, i.e., a variably heightened or lessened synchrony between the two groups, while there are no within-group changes. In the case that all coefficients of the projection vector are of the same sign, the correlations between brain regions of the 4-7 Hz periodic LFP components change in a more uniform way to an extent proportional to the product of their coefficients in the projection vector, i.e., a variably heightened or lessened synchrony across the 8 brain regions. The projection then comprises the direction of change in each brain region’s LFPs along which the coherency structure of the LFPs sustains a change after induction by CGRP.

#### Changepoint Clustering

For each possible combination pair of mice, we calculated the cosine distance between their projection directions. We then used this distance for k-medoids clustering. The number of clusters is estimated by optimum average silhouette width. And the clustering procedure is implemented on each of the three days for each frequency band.

## RESULTS

### ELS exacerbates migraine-related behavioral phenotypes in CD1 mice

Stress, especially stress experienced early in life, is a naturally-occurring event in the life of both humans and rodents that has been demonstrated to play an important role in migraine and has been implicated as altering plasticity in several affect-related brain regions including: the anterior cingulate cortex (ACC), amygdala (AMY), posterior thalamus (Po), ventroposteromedial thalamus (VPM), medial dorsal thalamus (MDthal), and parabrachial nucleus (PBN). ^10,11,70,71^. Specifically, adverse childhood events, also referred to as early life stress (ELS), are associated with migraine diagnosis in adulthood ^6,7^. In preclinical models, previous studies have found that ELS enhances migraine-like phenotypes in rats ^72^ and female C57Bl/6J mice ^73^, but importantly, this relationship hasn’t previously been identified in the context of CGRP.

We induced ELS in the outbred strain, CD1, mice by utilizing a maternal separation stress paradigm that also included early weaning and limited nesting (Figure 1A). A total of 52 litters were utilized for experiments, and animals were split into balanced groups according to weight and cohort prior to experiments (Figure 1A). Animals were randomly divided in an age-balanced way such that half of the litters underwent ELS and half remained group-housed (control). To assess the effect of ELS on migraine-related phenotypes, we administered peripheral PBS or CGRP to both groups (Control-Veh/Control-CGRP and ELS-Veh/ELS-CGRP). Following our stress paradigm (see Methods), we waited until mice reached adulthood (P66-69) and performed behavioral assays to assess anxiety-like and migraine-like behavior. We hypothesized that ELS would exacerbate migraine-related phenotypes in mice treated with CGRP compared to control mice.

**Figure 1.**
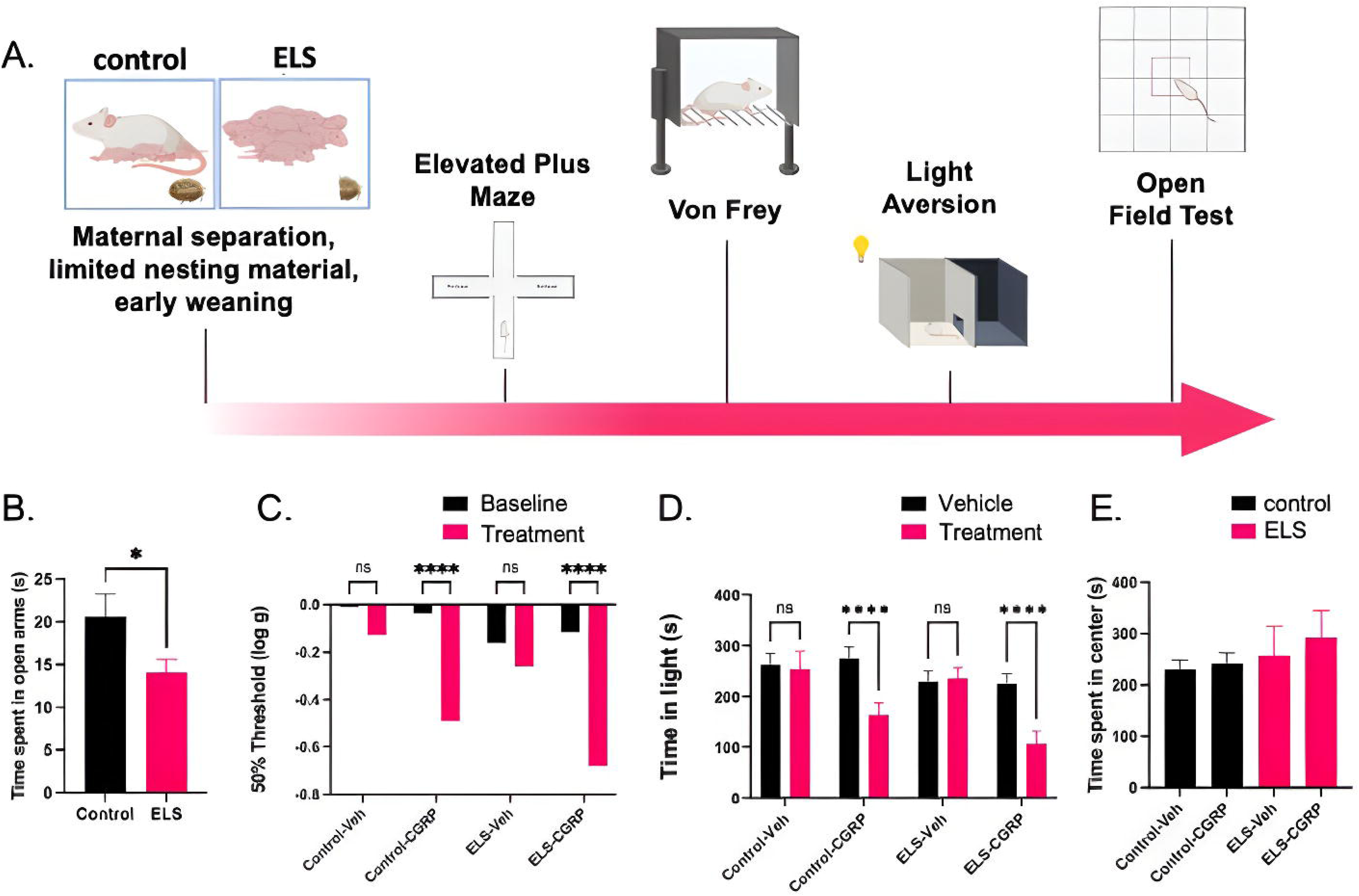
ELS exacerbates migraine-related behavior. A) Timeline of ELS experiment with behavior and recordings. (n= Control Veh:24, Control-CGRP: 24, ELS-Veh: 28, ELS-CGRP: 27) B) Animals spend less time in the open arms of the elevated plus maze following ELS C) CGRP-treated animals exhibit increased 50% threshold compared to baseline. D) CGRP-treated animals spend less time in the light during the light/dark assay. E) There were no significant differences in time spent in the center of the open field test following the light/dark assay.

First, we assessed anxiety-like behavior at P66-69 using the elevated plus maze and determined that animals exhibited an anxiety-like phenotype after undergoing ELS. We found that mice that underwent ELS spent less time in the open arms of the elevated plus maze (p = 0.04, t(91) = 2.11, unpaired t test) which indicates that the ELS stressed the animals sufficiently to demonstrate an affect-related phenotype in adulthood (Figure 1B).

Cutaneous allodynia is present in 60% of patients with migraine ^74^. We assessed cutaneous allodynia in CD1 mice at P72-79 by measuring mechanical sensitivity on the plantar surface of the hindpaw using the von Frey test. We log-transformed the data and utilized a mixed-effects analysis and found the interaction between ELS and CGRP treatment was significant (p<0.001). Using a Bonferroni’s multiple comparisons post-hoc test, we found that both Control-CGRP (p <0.001, t(94) = 5.92) and ELS-CGRP treated mice (p<0.001, t(94) = 8.18) exhibited increased mechanical sensitivity (50% threshold) compared to baseline sensitivity thresholds (Figure 1C). As expected, neither the Control-Veh (p =0.08, t(94) = 1.80) nor ELS-Veh (p = 0.16, t(94) = 1.41) groups exhibited significant differences in sensitivity thresholds.

At P81-85, we injected CGRP or PBS 30 minutes prior to assessing light-aversive behavior. Light aversion, or sensitivity to light, is present in nearly all patients with migraine, and about half of patients reported photophobia (light aversion) as the most bothersome symptom of migraine ^75^. The light/dark assay has been utilized to assess migraine-like behavior during the CGRP mouse model of migraine ^41^. Light aversion was measured by the average amount of time spent in the light zone of the light/dark box during the first 15 minutes of the testing period (Figure 1D). Using a Two-way ANOVA followed by Bonferroni’s multiple comparisons post-hoc test, we found that the interaction between ELS and CGRP treatment was significant (p<0.0001, F(3, 93) = 7.90). CGRP-treated control animals spent less time in the light following CGRP compared to PBS (p<0.001, t (93) = 4.55). CGRP-treated control animals also spent less time in light following CGRP compared to PBS (p<0.001, t(93) = 5.18). As expected, both Control-Veh (p = 0.74, t(93) = 0.337 and ELS-Veh (p = 0.78, t(93) = 0.283) did not show significant differences in time spent in light. Because the light/dark test can also be used to assess anxiety-like behavior, the standard in the field is to run an open field test under the same conditions as the light/dark test and verify that animals do not spend more time in center ^76^. We tested the animals at P83-86. We found that there were no significant differences in time spent in the center during the open field test (p= 0.76, F(3, 97) = 0.394) (Figure 1E), thus leading us to conclude that CGRP-treated animals did in fact demonstrate light aversion not attributable to anxiety during the test. Overall, we found that ELS predisposes CD1 mice to enhanced peripheral CGRP-induced migraine-related phenotypes.

### Peripheral injection of CGRP induces migraine-like behavior in electrode-implanted animals

To gain insight into brain mechanisms of migraine, and the ways that ELS impacts such mechanisms, we implanted electrodes and recorded neurophysiological activity from multi-site electrode wires in the ACC, AMY, MDthal, Po, VPM, and LPBN/RPBN (Figure 2A, Supplementary Figure S1). Such an approach allows us to quantify spatiotemporal dynamics following peripheral CGRP administration based on local field potential (LFP) activity and subsequently perform behavioral studies to phenotypically validate the effect of peripheral CGRP (Figure 2B). We first determined that our implanted animals exhibit similar light aversive behavior following CGRP administration as unimplanted animals (Supplemental Figure S2). There was no significant interaction between CGRP treatment and the electrode-implanted condition (p = 0.79). However, as expected, CGRP treatment did result in decreased time spent in the light on the CGRP treatment day compared to the vehicle day (p = 0.01, Supplemental Figure S2). We found no significant effect of repeated injection (p = 0.67, Supplemental Figure S2), nor due to the day itself (p = 0.77). Overall, we demonstrated that our electrode-implanted animals exhibit the same phenotype as unimplanted animals, consistent with phenotypes previously published ^42^. We also analyzed the total amount of time spent in light and resting time in dark across all implanted animals utilized in subsequent experiments (Figure S3). Again, implanted animals injected with CGRP exhibited decreased time spent in light during all 30 minutes (n = 31, p <0.001, Figure S2) and the first 15 minutes of the light/dark test (n = 31, p<0.001, Figure S2).

**Figure 2.**
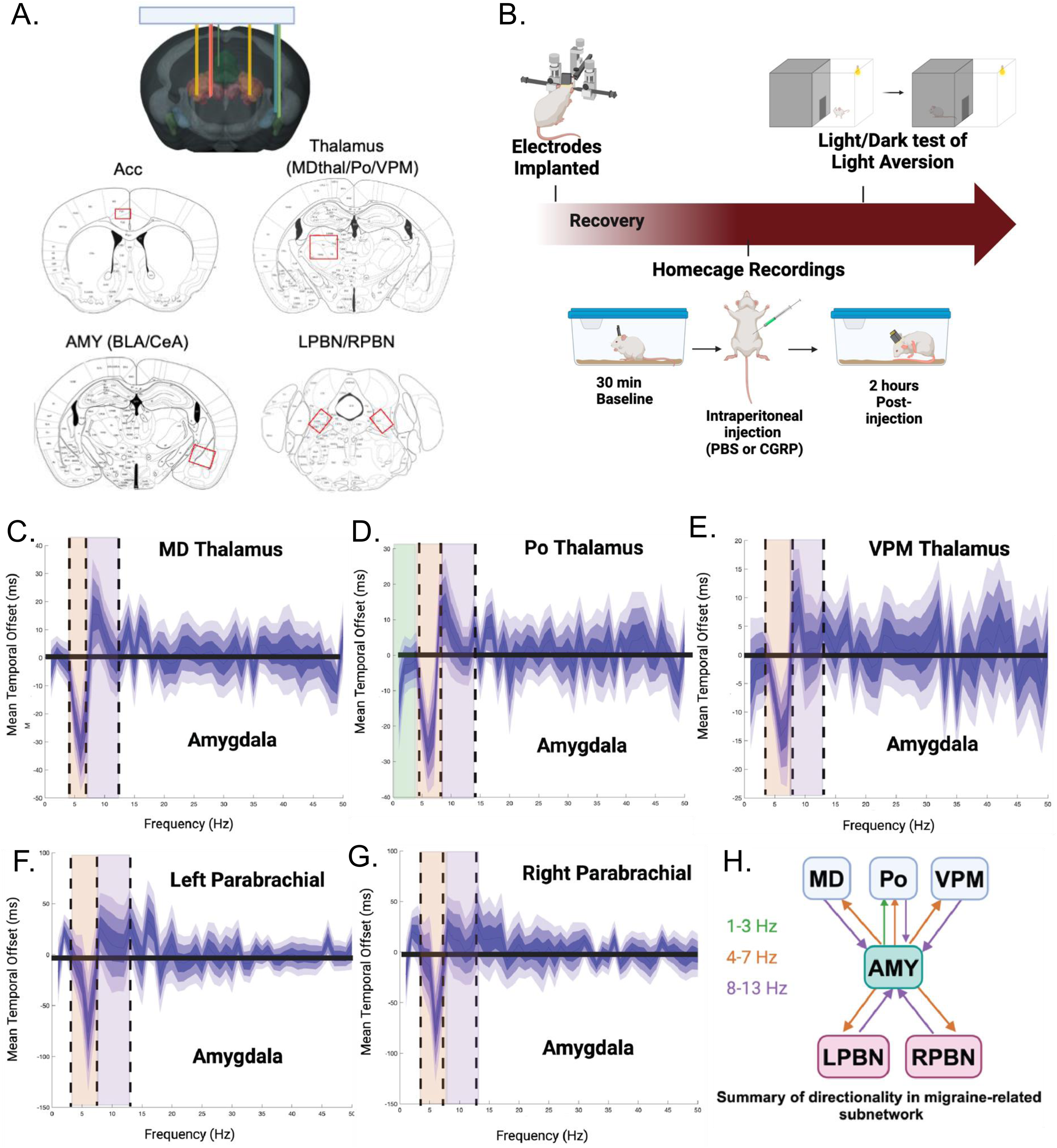
Multi-site in vivo Neurophysiological recordings identify AMY-centric network. A) Brain regions targeted in 32 CD1 mice (15 M, 17 F): ACC, AMY, VPM, MDthal, Po, and bilateral PBN (see also Figure S1). B) Experimental timeline of recordings and behavior C-H) LFP directional timing between brain region pairings across migraine-related subnetwork. The mean temporal offset at which each pair of LFPs optimally phase synchronized is plotted for each frequency 1-50 Hz. The solid blue lines indicated the mean temporal offset where lighter shaded blue represents standard deviations from the mean (1-3 standard deviations). C) MD– AMY D) Po–AMY E) VPM–AMY F) LPBN–AMY G) RPBN–AMY H) Summary: The mean temporal offset across the MD, Po, VPM, AMY, and PBN pairings are the greatest in the 1-3 Hz (green), AMY leads Po; 4-7 Hz (orange), AMY leads all components of the network; and 8-13 Hz (purple), all other regions lead AMY; frequency bands are indicated by the dashed lines. F) Summarized directionality relationships across all regions. Overall, LFP directionality in a migraine-related subnetwork is the greatest at the 4-7 and 8-13 Hz frequency bands.

### LFP directionality points to Amygdala-centric subnetwork

To identify LFP oscillation frequencies of interest, we calculated directionality of information flow from one brain region to another between brain region pairs based on a temporal lag approach in which phase offsets are introduced. Multiple studies have demonstrated important biology and behavioral predictions in frequencies where one region of a circuit clearly leads another between coherent brain regions ^57,65,77-81^. The temporal offset at maximal synchrony indicates which region is leading and lagging at each frequency ^57,79,82-84^. In order to specify precise frequency bands in which information flow appears to be directional from one region to another, we applied a fused lasso approach, enforcing a lasso penalty on absolute value differences between contiguous frequencies in phase offset (see methods, Figure 2).

Given the roles of the amygdala and thalamic nuclei in affect and migraine, and that they appear to play a hub role in migraine neural circuitry (Figure 8A), we focused on the directional relationships between these regions. To determine frequency bands of interest, we utilized a lasso regression on the greatest mean temporal phase offsets within these pairings (Figure 2). We found that AMY leads all of the thalamic (MD/Po/VPM) and bilateral parabrachial (LPBN/RPBN) nuclei in the 4-7 Hz frequency range (Figure 2). Continuing in this subnetwork, we found that all other regions led the AMY in the 8-13 Hz range (Figure 2). As such, we focused our migraine mechanistic studies on these frequency bands.

### CGRP induces decreases in LFP power across migraine-related brain regions

Our analysis indicates that 4-7 Hz is a frequency band in which the AMY is a major leading region. This frequency range has been identified in various brain regions to be involved in functions like sleep, memory, and anxiety ^83,85-90^. We hypothesized that we would observe significant differences in LFP power across each of the brain regions we implanted in the 4-7 Hz frequency. All data were normalized to the baseline period before plotting, and the time series was evaluated starting from the 30-minute mark, corresponding to the injection time for each animal. Because we’re interested in both the magnitude of response as well as the time-dependence, we employed a Function on Scalar General Additive Model analysis (FOS) followed by an FDR correction, testing the hypothesis that there are differences in LFP power across time in response to peripheral CGRP within each brain region. This analysis revealed that LFP power in the 4-7 Hz frequency band was significantly different across all regions we implanted over time when animals were treated with CGRP compared to when treated with PBS (Figure 3). Across all seven brain regions (ACC, AMY, MDthal, Po,VPM, LPBN, and RPBN) there was a decrease in LFP mean logpower following CGRP administration compared to vehicle control initially following the injection point (Figure 3A) and gradually increasing until reaching a similar mean logpower under the vehicle condition (Figure 3B-H). ACC (Figure 3B) contained the greatest relative changes immediately post-injection. The peak changes in power induced by CGRP occurred around 10-30 minutes post-injection, with the effects primarily resolving by 45 minutes post-injection (Figure 3, Supplemental Figures S6-7). Some regions leveled off in change by 60-min post injection or beyond but do not reach zero difference between CGRP and vehicle within the timeframe observed (e.g. Fig 3B,F,H, Supplemental Figures S6-7). The general timeframe of activity following peripheral CGRP injection from these data are highly consistent with what has been reported in terms of behavioral response to peripheral CGRP using facial grimace^66^. Notably, the VPM appears to be impacted first and resolve first in both frequency ranges, with the PBNs peaking and resolving on the later side (Supplemental Figures S6-7). ACC also was the last or among the last to resolve at both frequencies (Supplemental Figures S6-7).

**Figure 3.**
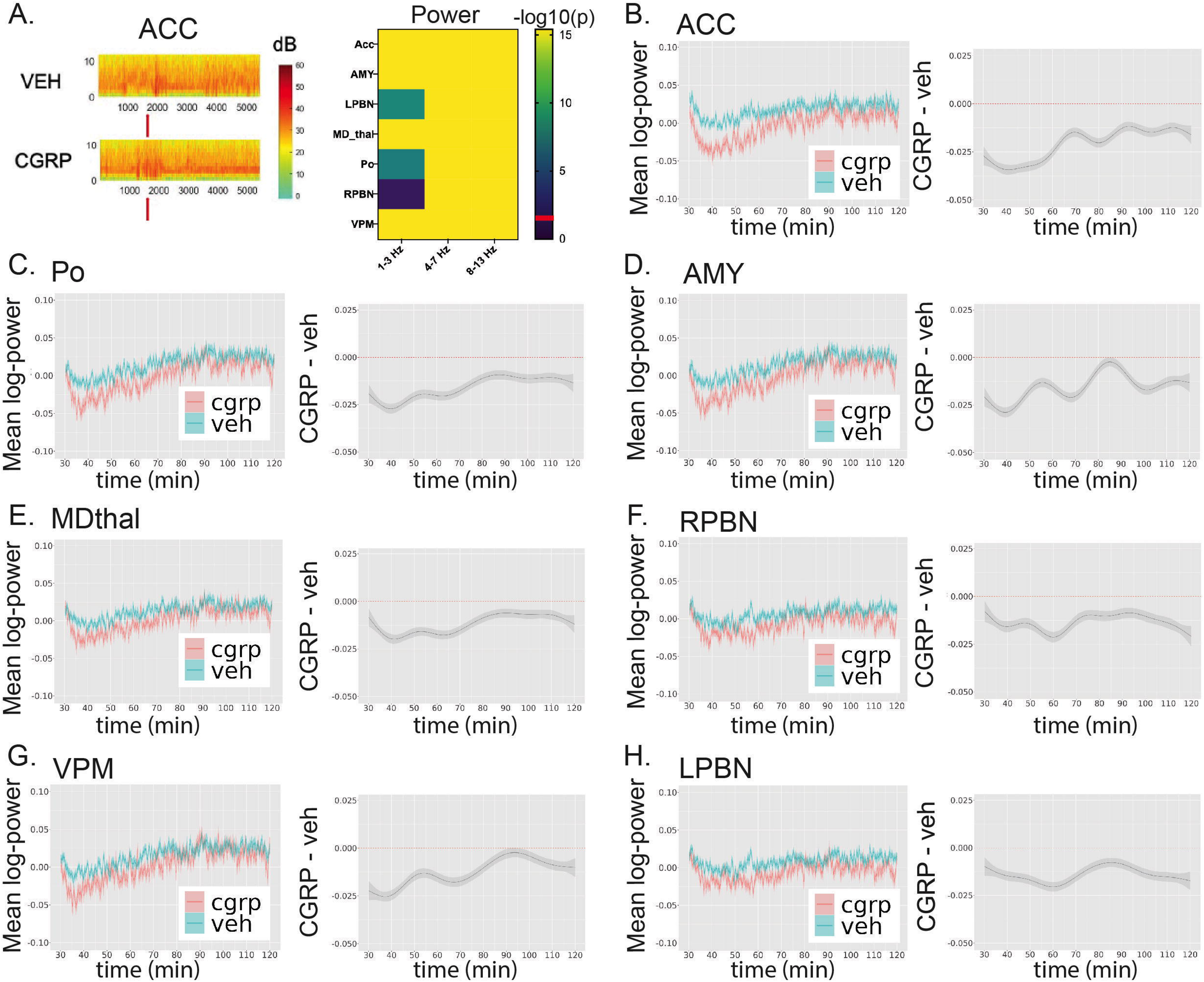
CGRP alters LFP power in the 4-7 Hz frequency band. LFP mean logpower plotted across time (n=32, see also Figure S3-4). A) Representative spectral power plot of the ACC across 1-13 Hz. Power is represented in decibels (dB) (left). Heat map showing all p-values across brain regions of interest (right). B-H) Plots showing the mean log-power across time following CGRP treatment (pink) compared to vehicle PBS (blue) (left) for each brain region. Function on scalar generalized additive model analysis (FOS) plots showing the difference between the mean log-power for CGRP and vehicle conditions immediately post-injection for each brain region (right). Data are represented as 30-second rolling means ± SEM.

We also hypothesized that the 8-13 Hz frequency band was a relevant frequency band as all of the other brain regions led the AMY in the 8-13 Hz frequency band (Figure 2). Similar to the 4-7 Hz frequency band, we observed changes in mean logpower in all regions of our network, though minimal in the 1-3 Hz band (Supplemental Figures S4-5). All three frequency bands were taken into account for our FDR corrections (see methods). In the 8-13 Hz band, the ACC (Supplemental Figure S5) contained the greatest relative difference, followed by the AMY and LPBN and overall power changes resolved by around 45-60 minutes post-injection (Supplemental Figure S5). Overall, we observed changes in LFP power across all brain regions of the network in two different frequency bands in response to peripheral CGRP, supporting centrally-mediated activity during a peripherally-induced migraine brain state.

### CGRP induces disruptions in LFP coherence across migraine-related brain regions

Next, we wanted to evaluate how functional connectivity across the network was altered, if at all, during a CGRP-induced migraine brain state, especially between AMY-brain region pairings. To answer the question of how brain regions communicate with each other across time during a migraine-like brain state, we analyzed coherence among brain region pairings following CGRP administration compared to vehicle conditions.

High mean coherence was observed across the major pairings of our network (e.g. pairings with AMY and Po hubs) across all frequencies of interest (i.e. 1-13 Hz) +/-CGRP (Figure 4A). All LFP data was normalized to baseline periods and plotted starting at the 30-minute point onward providing a stabilizing transform while preserving direction, as log odds coherence. Similar to CGRP-induced changes in power, the disruptions in coherence in response to CGRP were most pronounced in the first 10 minutes post-injection, but unlike the power, they largely resolved much faster, within ∼20 minutes post-injection (Figure 4B). The FOS model demonstrated that there are significant disruptions in coherence across network brain region pairings in response to CGRP compared to vehicle (Figure 4C).

**Figure 4.**
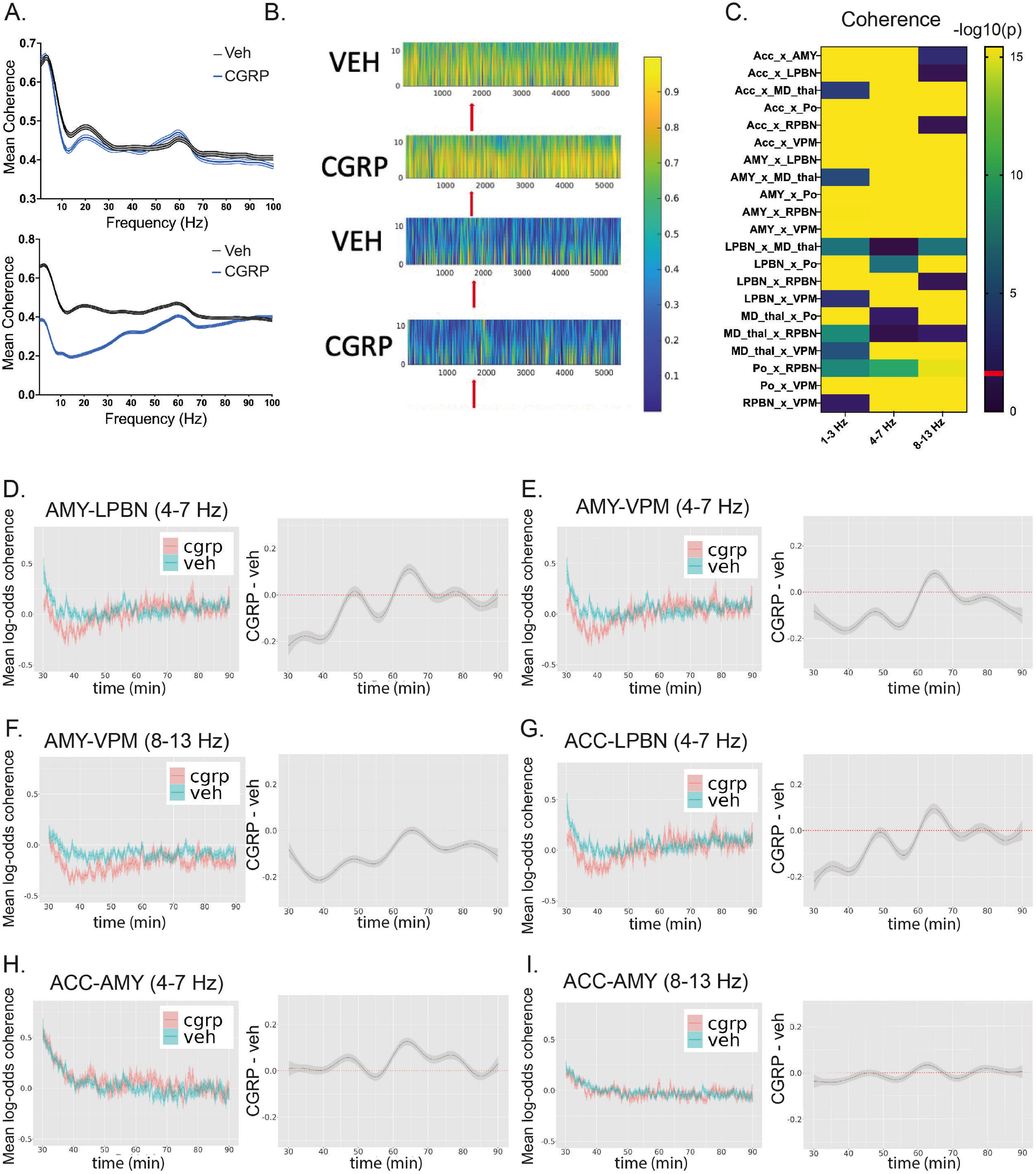
CGRP induces disruptions in LFP coherence across migraine-related brain regions. A) High mean coherence across all animals was observed during the 1-13 Hz frequency range (top) AMY-Po (bottom) AMY LPBN. B) Representative coherence plots of the (top) AMY-Po and (bottom) AMY-LPBN across 1-13 Hz. Red arrow indicates injection point. C) Heat map of p-values for differences between CGRP and vehicle mean log-odds coherence. D-F) For each brain region pairing, LFP mean log-odds coherence plotted across time for vehicle and CGRP conditions (left), and difference between vehicle and CGRP conditions (right) (see also Figure S5-6). Data are represented as 30-second rolling means ± SEM.

In both the 4-7 Hz band and 8-13 Hz frequency band, we observed significant coherence disruption in all AMY brain region pairings, the most of any of the regions (Figure 4C). Brain regions in which we observed the greatest magnitudes of CGRP-induced change in coherence (i.e. difference in mean log-odds coherence) were: the AMY-LPBN (0.218), AMY-VPM (0.213), and ACC-LPBN (0.223; all p= 3.74e-16, Figure 4D-G, Supplemental Figure S8-9). Notably, AMY-VPM coherence exhibited the most temporally sustained disruptions across time (∼35 minutes post-injection, p= 3.74e-16, Figure 4E-F). These findings support the critical role of electrical activity alterations in the amygdala during migraine attacks. Disruption of AMY-related LFP patterns appear to underlie the endogenous migraine brain state, leading to the painful and affective symptoms associated with migraine.

### Sumatriptan partially reverses the effect of CGRP

To contextualize the clinical relevance of these CGRP-induce brain network dynamics, we evaluated how CGRP-induced migraine-related brain network dynamics respond to a common acute migraine treatment, sumatriptan^91^. Sumatriptan is a serotonin 1B/D agonist that was originally thought to have peripheral mechanisms of action but recently has been found to act in the brain as well^92,93^. Brain network dynamics measured by fMRI of thalamus and amygdala have been demonstrated to be altered in response to sumatriptan ^94^. In other rodent models sumatriptan has had somewhat limited efficacy but does at least partially reduce CGRP-induced migraine-like phenotypes ^42,66^. We hypothesized that coadministration of sumatriptan and CGRP would attenuate some or all of the changes in power and coherence induced by CGRP.

We compared responses between CGRP alone and sumatriptan+CGRP and included vehicle and sumatriptan-alone conditions to assess the direction of the sumatriptan+CGRP relative to CGRP and vehicle (Figure 5, Supplemental Figures S10-11). Interestingly, ACC, and AMY in the 8-13 Hz frequency band were the only power features in which co-administration of sumatriptan+CGRP attenuated the CGRP response, with AMY demonstrating the most clear response (Figure 5B-C, p=3.74e-16). With regard to coherence in the 20 minutes following injection most disrupted by CGRP, we found only modest effects of sumatriptan+CGRP that occurred late in the timeframe (Supplemental Figure S10). Interestingly, the brain region with the greatest number of pairings reaching statistical significance and the greatest effect of sumatriptan blunting CGRP response was ACC, including ACC-AMY 8-13 Hz (p= 1.62e-13, Figure 5F), ACC-VPM 4-7Hz (p=7.5e-7, Figure 5H), 8-13Hz (p=3.74e-16, Figure 5G), ACC-Po 8-13Hz (p=4.68e-8, Figure S11), ACC-RPBN 4-7Hz (p=1e-6, Figure S10-11).

**Figure 5.**
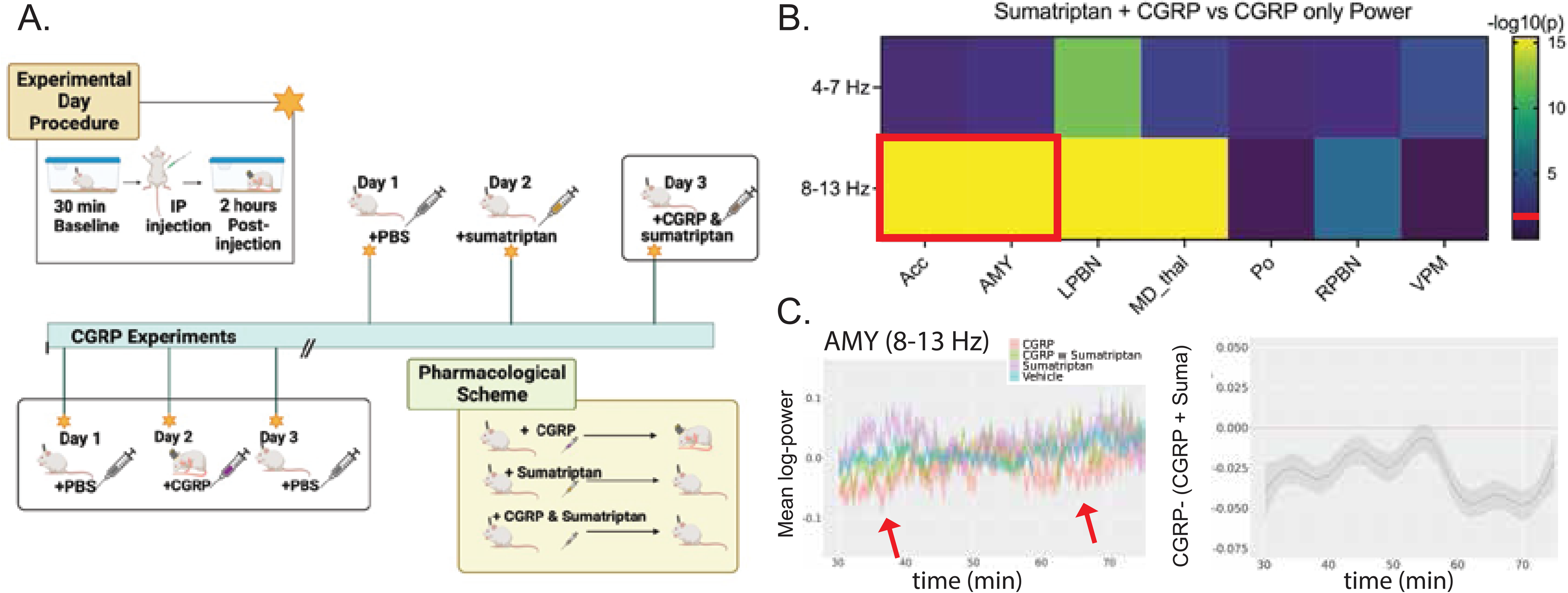
Coadministration of CGRP and Sumatriptan partially reverses the effect of CGRP. A) Experimental timeline of sumatriptan and CGRP experiments. B) Heat map showing -log10 p-values summarizing the effect of sumatriptan+CGRP on LFP power. Red boxes indicate significant differences in which reversal of the CGRP effect occurred. C) The mean log-power is plotted across time for the CGRP, sumatriptan+CGRP, and vehicle conditions (left) and difference between sumatriptan+CGRP and CGRP conditions (right). (top) ACC (bottom) AMY Power data are represented as 30-second rolling.

Sumatriptan also appeared to impact the network in additional ways beyond blunting the response to CGRP. The coherence changes between the LPBN and Po in the 8-13 Hz band, while significant, were not in the direction of CGRP reversal (p= 3.74e-16, Supplement Figure S10-11).

Overall, coadministration of sumatriptan and CGRP blunted the impact of CGRP on LFP power in the ACC, and AMY, and coherence across much of the network, especially in ACC-related region pairings and at later time points where the impact of CGRP is beginning to wane. These findings appear largely consistent with the timing and degree of sumatriptan blunting of CGRP in behavioral studies in mice^42,66^.

### ELS induces changes in LFP power and coherence in response to CGRP

Having identified a peripheral CGRP-induced migraine brain network response, we next set out to determine whether such network components could explain the exacerbating impact of ELS on CGRP-induced migraine-like phenotypes. Given the role of the AMY in stress and affect and the changes in LFP power and coherence we observed in response to CGRP, we hypothesized that animals exposed to ELS would exhibit significant differences in LFP power in the AMY and AMY-related coherence in response to CGRP, compared to animals that remained group-housed during early life. ELS and group-housed control animals were treated with vehicle on Days 1 and 3, and CGRP on Day 2. Overall, the ELS-CGRP exhibited the strongest decrease from baseline of relative power and coherence across regions and pairings and the greatest amount of changes from baseline compared to control groups (ELS-vehicle, Control-vehicle, Control-CGRP, Figure 6). We examined whether there was a significant interaction between ELS and CGRP response across time in these groups, as well as network-wide impacts of ELS itself in the absence of CGRP. We found significant interaction between ELS and CGRP with regard to LFP power in either the 4-7 Hz or 8-13 Hz to varying degrees across all brain regions of the network (i.e., a significant interaction, Figure 6, Supplemental Figure S-9).

**Figure 6.**
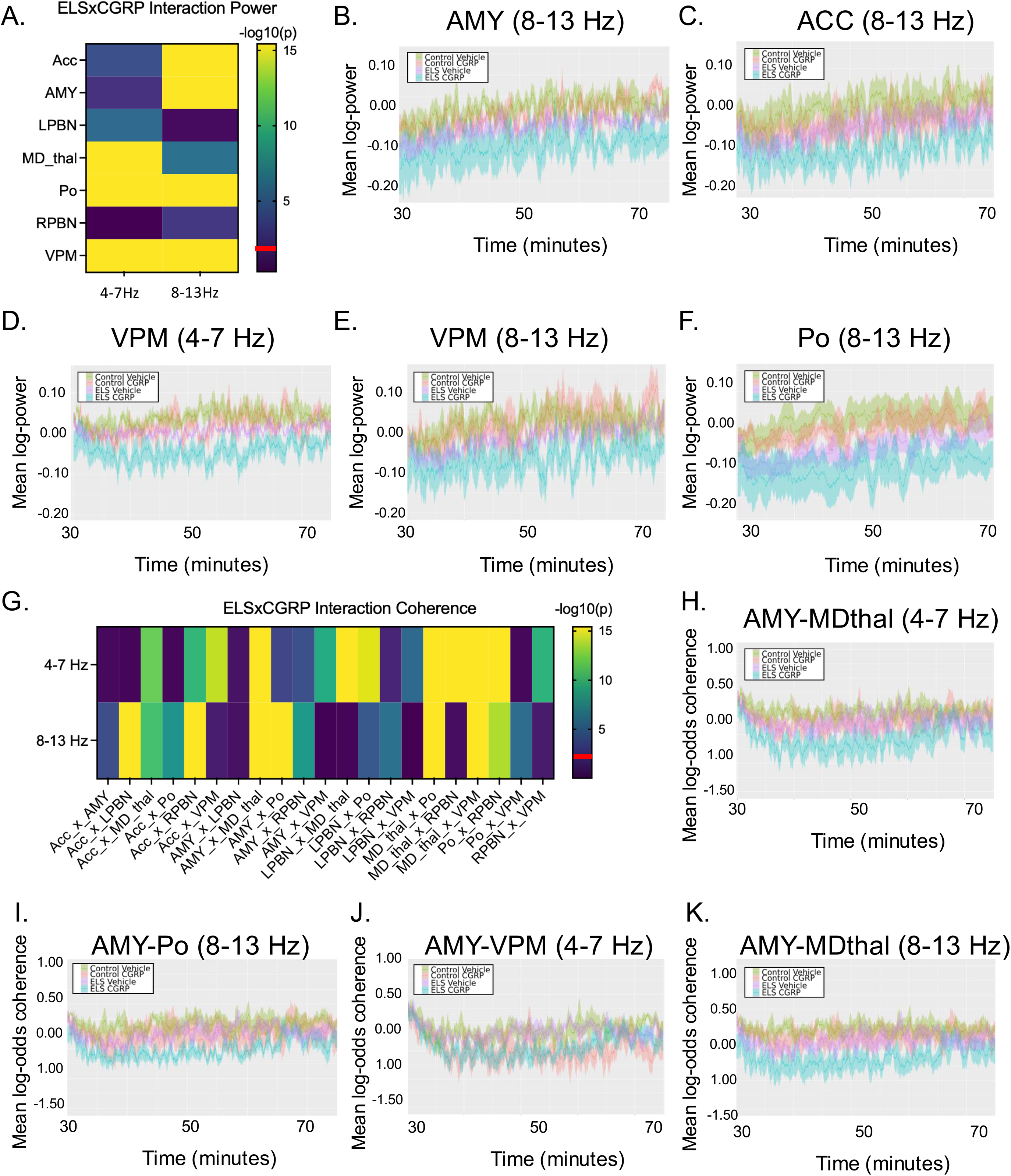
ELS alters AMY and Po-related circuitry in the presence of CGRP. A) Heat map of -log10 p-values reflecting the interaction between ELS x CGRP in ELS on LFP power (see also Figure S10) (n=11), B-G) the mean logpower in the AMY and Po are plotted across time for CGRP and vehicle conditions in ELS and control mice. Power data are represented as 30-second rolling means mean ± SEM. I) Heat map of -log10 p-values reflecting the interaction between ELS x CGRP in ELS on LFP coherence J-L) Mean log-odds coherence is plotted across time for Po-related region pairings for CGRP and vehicle conditions in ELS and control mice (see also Figure S12-13). Coherence data are represented as 60-second rolling means ± SEM.

With regard to ELS x CGRP interaction in power, in the 8-13Hz band, ACC, AMY, Po and VPM were the most significant features (Figure 6), whereas in the 4-7Hz range, the thalamic regions were (Po, VPM, MDthal, Figure 6, S12-13), implying likely AMY x thalamus mechanisms. There were no significant effects of ELS alone in any brain region or frequency (Supplemental Figure S14).

Given the differences in power reactivities during the ELSxCGRP interaction, we examined whether coherence patterns in the network were also impacted (Figure 6G). Many significant differences in coherence were observed between control and ELS groups after CGRP administration, including AMY-related pairings, which were consistently altered. Specifically, coherence was most disrupted in the ELS group in response to CGRP for AMY-thalamus pairings; AMY-Po 8-13Hz (ELS x CGRP interaction p= 3.74e-16, ELS effect p= 0.79, Figure 6I) and AMY-MDthal 4-7Hz, 8-13Hz (ELS x CGRP interaction p= 3.74e-16, ELS effect p= 0.68 Figure 6H, K) pairings, and AMY-VPM 4-7 Hz (interaction p=3.57e-10, ELS effect p=0.86, Figure 6J). When considering coherence pairings that had been most-impacted by CGRP, beyond AMY-VPM, AMY-LPBN showed no significant interaction of ELS and CGRP (4-7 Hz p=0.18; 8-13Hz p=0.22), ACC-LPBN demonstrated no interaction effect at 4-7Hz (p=0.66) but a strong interaction at 8-13 Hz (3.74e-16), and LPBN-Po demonstrated a moderate interaction (ELS x CGRP interaction p= 5.65e-5, ELS effect p= 0.67, Supplemental Figure S13).

Overall, these findings indicate that ELS induces changes across the network, and especially AMY and thalamic-related brain circuits in the context of CGRP across critical neural features that are responsive to CGRP. Given that there is no effect of ELS alone on any of these features suggests that ELS sensitizes the brain’s responsiveness to CGRP. Several coherence pairings that are most responsive to CGRP (AMY-Po and AMY-VPM) are rendered even more responsive with pretreatment with ELS. Our data thus demonstrate that ELS predisposes mice to a CGRP-susceptible brain state, resulting in compromised brain circuitry within CGRP-related pain pathways in the presence of CGRP. Such findings establish a novel model that predisposes rodents to a migraine-susceptible state (behaviorally and neurophysiologically), enabling rigorous investigation of dysfunctional migraine-related circuitry and providing a framework for studying the CGRP-vulnerable brain state (Figure 8).

### Changepoint Analysis identifies differing brain network responses to peripherally delivered CGRP

Up to this point, we have provided a thorough demonstration of the average changes in brain activity across a population of mice over time during peripheral induction of migraine with CGRP and the ways in which ELS impacts these network components toward a vulnerability to CGRP. In order to investigate the possibility of individual vulnerability to CGRP, we employed a change-point analysis to examine the evolution of important network-wide events in each individual in the CGRP-induction of a migraine-like state. This spectral-based method is used for detecting changes in the correlation structure of specific frequency components in the LFP (time series), over time, and the “principal component” of the LFPs, along which a change occurs. The change-point detection identifies the epochs (i.e. the change points) across which a data-driven linear combination of the periodic components of LFP in our frequency bands of interest undergo a variance change. Thus, a changepoint reflects a point where coherence across multiple brain regions have a coordinated and significant change. We reasoned that the first changepoint after the injection is likely to be the most informative, as this likely reflects an animal’s initial CNS response to the peripheral CGRP. The first changepoint after the injection (at 30 min), occurred within 40 minutes post-injection across all three days with no obvious relationship between light aversion behaviors and the timing of the first changepoint (Figure 7A). It does seem, however, that the time intervals between the change point and injection are smaller on day 2, the day CGRP was injected, than the vehicle days, 1 and 3. In order to ensure robustness of these findings, an additional robustness step was implemented, where we repeated the change point detection algorithm 50 times with different initialization random seeds, and evaluated the output change point locations by the mode and the projections by the median.

**Figure 7.**
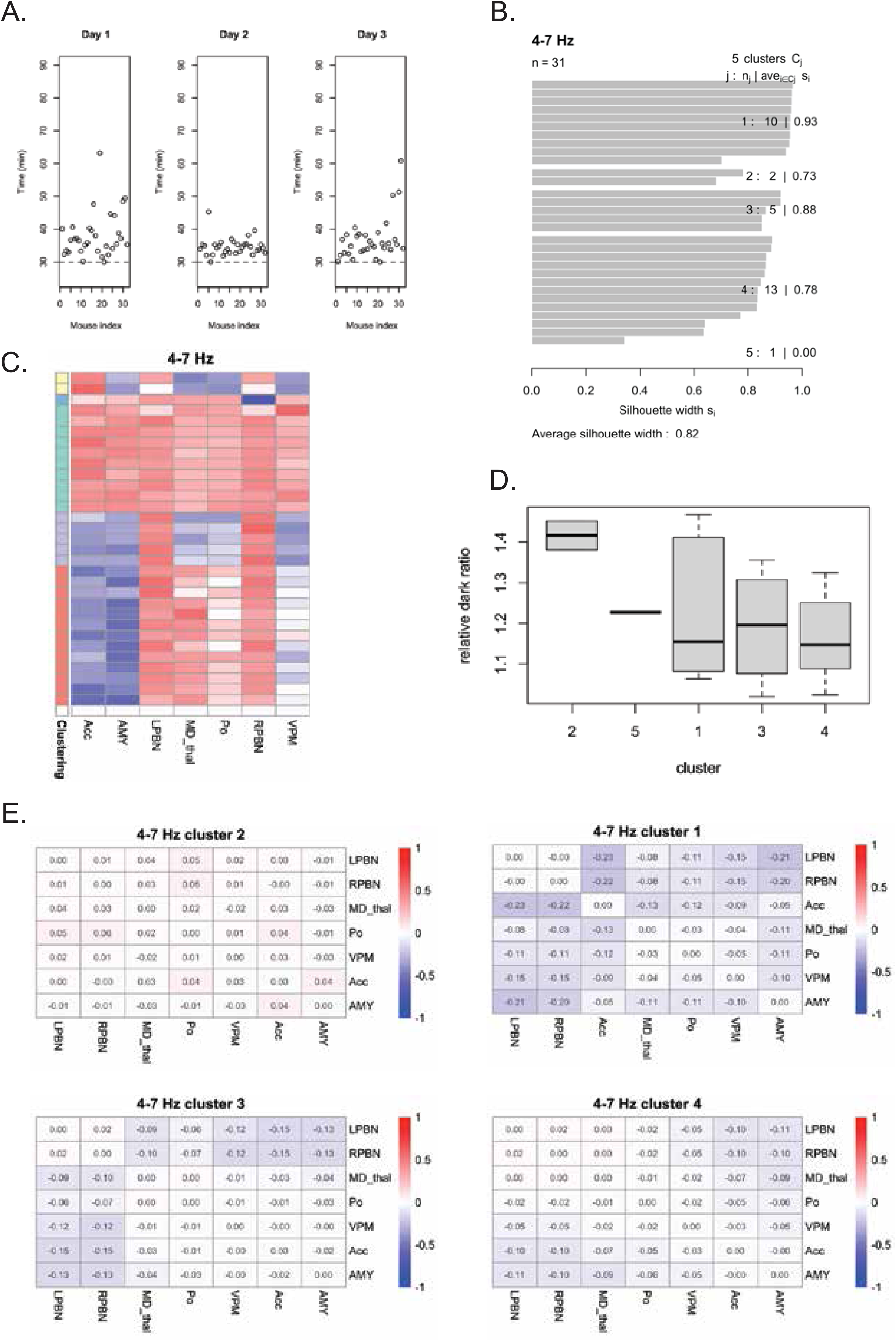
Changepoint Analysis identifies differing brain network responses to peripherally delivered CGRP. A) Time of 1^st^ change point after injection (at 30min) is shown on the y-axis, with mice shown in decreasing order of their behavior from greatest to least photophobia following CGRP. B) 4-7Hz k-medoids clustering of initial changepoint projection features following injection with CGRP, silhouette width on the x axis, and each mouse on the y axis, the counts and average silhouette width in each cluster listed on the right, and the overall average silhouette width on the bottom. C) 4-7Hz projection features for each mouse by cluster (colors to left). Brain regions are represented across the bottom with each mouse representing a different row, ordered by the cluster average of highest relative dark ratio (most photophobic) at the top. The legend shows the color of values of each cluster in the heatmap. D) light-aversive behavior (y-axis) for 4-7 Hz clusters demonstrate relationship between brain network feature clustering and behavior on the light/dark test, represented by the relative dark ratio= Treatment time in dark/ Vehicle time in dark. E) The cluster mean of difference of coherency matrix for 4-7 Hz over the first change point (i.e. the change in coherency matrices from the segment preceding changepoint to the segment after changepoint) after injection (in clusters with at least 2 mice and ordered by the average relative dark ratio) (see also Figure S15-18).

For a given changepoint, there exists a coefficient “projection” vector describing the brain region relationships that characterize the changepoint, which could be due to an altered synchronization strength among those brain regions in a given frequency band (see Fig. 7 and below). In order to determine whether there are common features regarding connectivity changes at the changepoint across animals in response to CGRP, we conducted a clustering analysis of the mice. We clustered in an unsupervised way based on the LFP features involved in each animals’ changepoint, by using a cosine distance between the projection vectors of the mice. We reasoned that different CD1 mice might respond differently at the level of the CNS in response to peripheral CGRP (i.e. have different projections at the first change point) and we therefore used a k-medoids clustering approach to identify groups of mice that respond with similar projections. We used a silhouette coefficient to assess the strength of the clustering structure with average silhouette width >0.8 for each frequency (4-7Hz and 8-13 Hz). We found four (4-7Hz) and three (8-13Hz) clusters this way, suggesting different brain network activity responses to CGRP in the CD1 mouse brain (Figure 7B, S15).

In order to understand the functional differences in the nature of changes in connectivity associated with these four different types of changepoints, we plotted the projection vectors for all of the animals grouped by cluster and ordered them from groups that are most to least photophobic in response to CGRP (Figure 7 C,D). As shown in Figure 7C, different coherency organizational structures, as indicated by the projection vectors (see Methods), define groups that are more or less susceptible to the migraine-inducing effects of peripheral CGRP injection. In other words, the network activity around the first changepoint explains the behavioral phenotype. To quantify the extent to which network activity can be used to predict hypersusceptibility to migraine in the mice, we calculated an AUC for the most highly susceptible group compared to the rest of the animals for 4-7Hz and 8-13Hz (AUC=0.92, p=0.028 4-7Hz; AUC=0.61, p=0.48, 8-13 Hz). In order to assess whether this strong relationship at 4-7Hz between brain network activity and behavior was specific for CGRP we did the same comparison for days 1 and 3 and found much weaker associations with behavior (day 1 AUC=0.72, p=0.06, day 3 AUC=0.68,p=0.24). Altogether, these data demonstrate that there is not one global response in connectivity of all animals to CGRP, that is to say that subgroups of different animals have differing connectivity response to peripheral CGRP injection across the brain. These data also demonstrate that coherence changes at the first timepoint following CGRP injection can be used to predict animals with extreme migraine-like responses based on 4-7HZ LFP activity.

In order to provide a more direct biological interpretation of the network features defining this network of migraine hypersusceptibility, we examined the coherency matrices immediately before and immediately following the changepoint (Supplemental Fig. S16-S17). In Figure 7E, we show the cluster mean of difference of coherency matrix on 4-7 Hz over the first change point after injection by cluster. For the 4-7 Hz frequency band, clear block-wise structure is observed for cluster 1 and 3. Interestingly, cluster 1, the cluster in which the animals have the strongest vulnerability to CGRP as measured by photophobia, is notably the cluster in which coherence is most increased, rather than decreased, across the network at the first changepoint following CGRP injection. This is consistent with the idea of the migraine response being concerted across multiple synchronized brain regions during specific events, while desynchronizing networks overall.

## DISCUSSION

Migraine is a complex and profound disorder, which includes experiences of pain and episodic disabling brain function, impacting up to 20% of the population at some point (∼1 billion people worldwide). Stress is a common trigger for migraine; high levels of stress are reported in migraine patients and, simultaneously, migraine is comorbid with numerous brain disorders associated with stress ^95,96^. In humans, early life adversity has a well-documented association with migraine and headache in adulthood^97^. Studies using rodent models have begun to investigate mechanisms by which this occurs ^98^. In this study, we sought to contribute substantially to our understanding of this mystery of how early life events can result in migraine symptoms in adulthood and found that neural activity within the amygdala and its functional connectivity network play important roles.

To understand the development of such a complex disorder as migraine, which induces systemic and coordinated malfunction of brain systems spanning emotion, sensation, autonomic nervous system regulation, and cognition, a systems neuroscience approach is needed ^39^. The field’s understanding of migraine has evolved significantly, shifting from thinking of it as a primarily neurovascular disorder to a complex neurological condition involving multiple brain regions, with different regions engaged during distinct phases ^99,100^. Several pieces of evidence point toward both central and peripheral pathophysiology, including the complex symptomatology experienced by patients with migraine, differences in functional connectivity between patients and healthy controls, and varied pharmacological responses; how exactly brain networks coordinate over time to produce a migraine brain state remain unknown. In this study, we have implemented a systems-based approach combining multi-site in vivo neurophysiology with peripheral CGRP injection, and migraine-relevant behavioral assays, in order to understand a peripherally induced migraine brain state, and the ways in which ELS impacts such a brain state.

We used peripheral injection of CGRP to best parallel human studies and account for the most universally understood mechanisms of how CGRP initiates a migraine episode. There are several plausible explanations of how peripheral CGRP ultimately regulates brain function, beginning with nociceptive primary afferents expressing and releasing CGRP providing input to second-order neurons of the spinal dorsal horn and trigeminal nucleus caudalis ^33,101^. This input is then transmitted to the parabrachial (PBN) and thalamic nuclei. A monosynaptic connection between trigeminal afferents and the PBN also exists, which can transduce craniofacial affective pain and exist in the dura ^37^ ^32^. CGRP is also expressed in projections from PBN neurons to the thalamus and amygdala ^102^. Additionally, CGRP can also directly or indirectly modulate neuronal activity and synaptic plasticity in several brain regions implicated in migraine, including the cortex, amygdala and thalamus ^102-104^. These data all contribute to our hypothesis that peripheral CGRP facilitates nociceptive transmission into the brain and contributes to the sensitization of both primary afferent sensory and second-order pain neurons to contribute to a migraine brain state.

Importantly, by identifying frequency bands with directionality to the flow of information (Figure 2), our results are interpretable in terms of AMY-leading (4-7Hz) and AMY-receiving (8-13 Hz) oscillations. Quantifying spatiotemporal dynamics following peripheral CGRP administration resulted in identification of coordinate changes in power and coherence, which are particularly interesting when considering the context of migraine as a phasic disorder. In the 4-7 Hz range, we found that CGRP injection led to a decrease in power across all 8 regions which peaked at 10-30 minutes post-injection, primarily resolved or leveled off by 45 minutes post-injection (Figure 3). A similar time course of CGRP-induced behavioral sensitization was found in a study where peripherally administered CGRP induced a significantly greater spontaneous facial pain response than controls starting 10 min post-injection, peaking at 30 minutes, and lasting for about 75-90 minutes ^66^. This is consistent with other preclinical mouse models of migraine, and, to some extent, human studies of photophobia ^42,105^. A recent fMRI study of patients across the prodrome to pain state transition also reported a large drop in widespread brain network connectivity from the transition between prodrome and the pain/headache attack phase as well^106^. While a lot of heterogeneity exists with regard to human fMRI studies broadly, globally, thalamic connectivity entering a headache attack is both strengthened and weakened, our findings are largely consistent with human studies of the migraine attack phase as a whole^100,107^. We observed that VPM was the first region to reach its maximal activity and the first region for CGRP-induced changes in LFP power to resolve. This suggests that VPM plays an important role in the initiation of the migraine response across the brain, perhaps initiating through projections it receives from SpV ^25,108^. The ACC was a region that was last to resolve, extending beyond the timeframe of most observed migraine-like behaviors^66^. Given the role of the ACC in processing the emotional aspects of pain, and that LTP has been found to be involved in the ACC in pain and migraine, this finding may point to a possible role for the ACC in post-drome or interictal migraine phases^109^. Imaging studies of the postdrome phase have understandably been quite limited in numbers of patients and brain regions examined^100,110^. Additionally, in light of the field’s ongoing debate about peripheral vs. central mechanisms of CGRP, and the fact that CGRP largely does not cross the blood-brain barrier, it is especially important that we have begun to map out specific activity across the brain in response to a peripheral inject of CGRP. Notably, the half-life of CGRP is ∼6 minutes and yet we see sustained activity across the brain, pointing to a peripherally triggered process that gets sustained at much higher levels of processing.

Given our overall network connectivity hypotheses, we were interested in whether CGRP disrupted coherence between AMY, ACC, and thalamic regions, and as these regions form key circuits involved in chronic pain ^111^ and pain perception ^112^. Coherence was disrupted across the network shortly after injection with CGRP and across the board followed a shorter time course than power disruptions. Significant disruptions in coherence among brain region pairings were especially prevalent in the first 10 minutes immediately post–injection of CGRP, and resolved around 20 minutes post-injection. This time course gives us insight into how information flow between brain regions is disrupted by a migraine trigger to orchestrate the migraine brain state. Because CGRP enhances sensitivity to sensory input in both the periphery and central nervous system, it is possible the changes in coherence are a reflection of a transition into a hypersensitized brain state ^113^.

As a well-established therapeutic for migraine, sumatriptan provided an important test of the impact of a clinically relevant perturbant on the CGRP-responsive network. Mechanistically, sumatriptan is part of a triptan class of 5-HT_1B/1D_ receptor agonists that can directly suppress migraine-related increases in CGRP secretion from trigeminal neurons and normalize cranial CGRP levels, however, the site of action is debated ^92,94,114-116^. It can prevent the induction of sensitization in central trigeminovascular neurons but also has recently been shown to reach brainstem and hypothalamic structures quite rapidly^92,117,118^. Sumatriptan has partially reversed behavioral phenotypes of peripheral CGRP injection previously^66^. Coadministration of CGRP and sumatriptan reversed the changes in power in the ACC and AMY in the 8-13 Hz frequency band especially at 25 minutes post-injection (Figure 5, Supplemental Figures S10-11). Sumatriptan reversed coherence disruptions in brain region pairings involving the AMY and ACC, especially within the first 20 minutes post-injection. An fMRI study with normal, healthy volunteers demonstrate that peripheral CGRP-induced decreases in BOLD-signal in the caudate nuclei, thalamus, and cingulate cortex were reversed by sumatriptan at 40 minutes post-CGRP injection and 15 minutes post-sumatriptan injection. ^119^. Our data seem to recapitulate at least the cingulate aspect of this finding in our mouse model. Given that sumatriptan doesn’t ablate all CGRP-induced effects and even introduces some additional connectivity patterns, such brain connectivity metrics may be useful to consider in therapeutic development moving forward with regard to optimizing network responsivity. Future studies should also investigate the effect of CGRP-targeting pharmacologics (e.g., CGRP monoclonal antibodies, CGRP antagonists) on CGRP-induced brain states to assess whether network activity is more strongly attenuated.

While other studies in rodent models have investigated a role for ELS in migraine, to the best of our knowledge, ELS had not yet been investigated in a CGRP mouse model of migraine, nor in CD1 mice. In our study, ELS significantly enhanced the CGRP-induced photophobia and allodynia (Figure 1). Further, we investigated the impact of ELS on brain network activity and found that ELS significantly enhanced the CGRP-induced drop in power found across the network, especially with regard to the amygdala. Coherence patterns demonstrated that most, but not all, of the network connectivity pairings that increase most with CGRP are significantly exacerbated by ELS (AMY-VPM, AMY-Po), providing a plausible mechanism by which ELS sensitizes the brain to CGRP response. Importantly, AMY-MDthal connectivity also displayed significantly increased CGRP response in ELS animals, which is especially interesting given the strong relationship between both of these regions and stress response in the context of mood disorders^120,121^. Globally, ELS appears to especially predispose the network so that CGRP drives amygdalar-thalamic coherence even further. Given the role of the thalamus in sensory gain/sensitivity and the role of the amygdala in the emotional aspect of pain, CGRP-dependent dysregulation of the functional connectivity between these two regions in the context of ELS appears to be an important aspect of the way that ELS impacts migraine in adulthood. Given that ELS alone has no significant impact on the network activity, the interaction of ELS and CGRP must derive from some network property that is not actively engaged until it gets unmasked upon stimulation with CGRP. In short, ELS predisposes mice to a CGRP-vulnerability brain state. Now that we have determined a way to induce and measure such a state, future studies can now more deeply investigate the detailed cellular and molecular contributors of this CGRP-vulnerable brain state (Figure 8).

**Figure 8.**
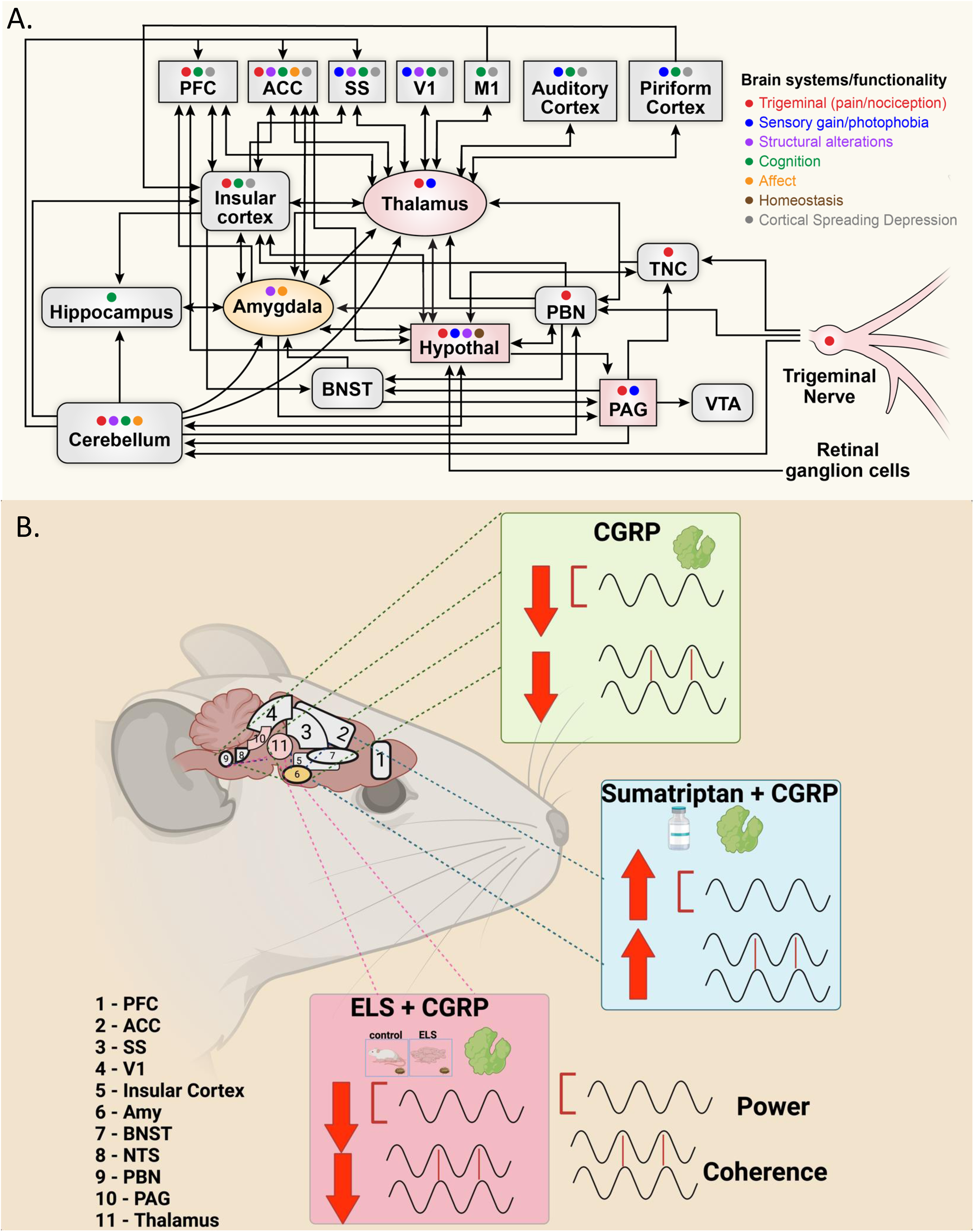
Model of Migraine Network Activity. Top: Brain network components implicated in migraine either by human fMRI or by rodent pain studies. *Abbreviations*: ACC-anterior cingulate cortex, Amy-amygdala, BNST-bed nucleus of the stria terminalis, Hypothal-hypothalamus, M1-motor cortex, NTS-nucleus tractus solitarius, PAG-periaqueductal gray, PFC-prefrontal cortex, PBN-parabrachial nucleus, SS-somatosensory cortex, TNC-trigeminal nucleus caudalis, V1-visual cortex, VTA-ventral tegmental area. Figure created by first organizing human migraine imaging literature to identify key brain regions involved in migraine and then organizing network connectivity by reviewing preclinical migraine and related circuits. Finally, circuit connections were filled in between the major hubs and network components. ^3,11,21,28,33,37-39,100,101,122-147^. Map is not meant to be comprehensive, especially between cortical regions. Also not shown are spinal inputs, which can directly impact thalamic and hypothalamic regions, among others ^25,148^. Bottom: Model of brain network activity in peripheral CGRP-induced migraine state.

These global findings provide an overall picture of brain network responses generalized across the population that we measured and ways in which this network is altered by ELS, however, our changepoint analysis best helps us understand what brain network activity might be most contributing to individualized migraine-like responses. Given the heterogeneity of migraine experience, we thought this was especially important to pursue. We identified changepoints, which are time epochs where LFP signals within specific frequency bands undergo a substantial and coordinated change in variance (Figure 7). Our change-point analysis clusters suggest different brain network activity responses to CGRP in the CD1 mouse brain. These clusters, when ordered from most to least photophobic in response to CGRP, revealed different coherency organizational structures which defined groups that are more or less susceptible to peripheral CGRP injection-induced migraine. Importantly, the network activity around the first changepoint – 10 minutes post-injection – provides sufficient information to explain or even to some extent predict, the behavioral phenotype. Thus, coherence changes at this first timepoint following CGRP injection can be used to predict animals with extreme migraine-like response to light. The cluster of animals with the most photophobic response to CGRP is the only cluster in which coherence in the 4-7 Hz frequency range is increased at the first changepoint, suggesting an important transient transition of brain network activity specifically in highly migraine susceptible animals that differs from the overall population. Pinpointing the differences between migraine susceptible and resilient brain network activity opens up exciting possibilities for studying migraine resilience and the ways in which environmental contributors such as ELS make some individual open to migraine vulnerability.

Our findings provide an important roadmap of endogenous brain network activity in response to arguably the most clinically relevant mouse model of migraine, induction by peripheral CGRP. In a context where central and peripheral mechanisms of CGRP’s influence in migraine have been debated for so long, this map with both global (multi-region connectivity) and neural timescale information provides the needed framework for the next era of work to understanding brain response to peripheral physiology. Moreover, we have demonstrated that such metrics can provide important answers about clinically relevant problems such as the ways in which ELS predisposes the brain to respond to CGRP. Our results provide unique insights into the neural basis of migraine and provide a reasonable launching point for the development of novel therapies targeting brain-wide networks and potential targets for neuromodulation.

## CONCLUSIONS

We describe a brain-wide network neural activity response to peripheral induction of migraine-like activity by CGRP. This is an important step forward in the field because CGRP does not cross the blood brain barrier and the degree to which peripheral induction of CGRP impacts brain activity has been an ongoing debate in the field. Furthermore, we provide evidence for a mechanism by which ELS predisposes the brain to respond to this common migraine trigger of peripheral CGRP. We anticipate that these findings open doors for future brain-network based therapeutic development for migraine.

## RESOURCE AVAILABILITY

### Lead contact

Further information and requests for resources and reagents should be directed to and will be fulfilled by the lead contact, Dr. Rainbo Hultman

### Materials availability

There were no materials generated during this study.

### Data and code availability

- All original code has been deposited at GitLab and will be publicly available as of the date of publication.
- Any additional information required to reanalyze the data reported in this paper is available from the lead contact upon request.

## OTHER ACKNOWLEDGMENTS

We would also like to thank Andy Russo and Levi Sowers for helpful conversations and pharmacological expertise and Kiriana Williams for technical assistance. We would like to acknowledge Biorender, which was used to make some of the figures.

## AUTHOR CONTRIBUTIONS

Conceptualization, R.H., S.S., and M.J. methodology, M.J., M.E., I.H., X.Z.,; Investigation, M.J. M.E.,Y.F.,; technical support, M.J., M.E.,Y.F., A.J., M.M., S.M., R.V., and J.M.; writing original draft, R.H., M.J., A.A., and M.E.; writing—review & editing, M.J., M.E., I.H., and X.Z.,; funding acquisition, R.H.. and S.S..; resources, I.H., X.Z., S.S.. and K.S.C..; supervision, R.H., S.S., B.H., and K.S.C.

## DECLARATION OF GENERATIVE AI AND AI-ASSISTED TECHNOLOGIES

The authors have nothing to declare regarding AI.

## DATA AVAILABILITY

*The data used in this article will be shared on reasonable request to the corresponding author*.

