## Supplemental Materials for "Early Life Stress induces brain-wide electrical network predisposition to migraine"

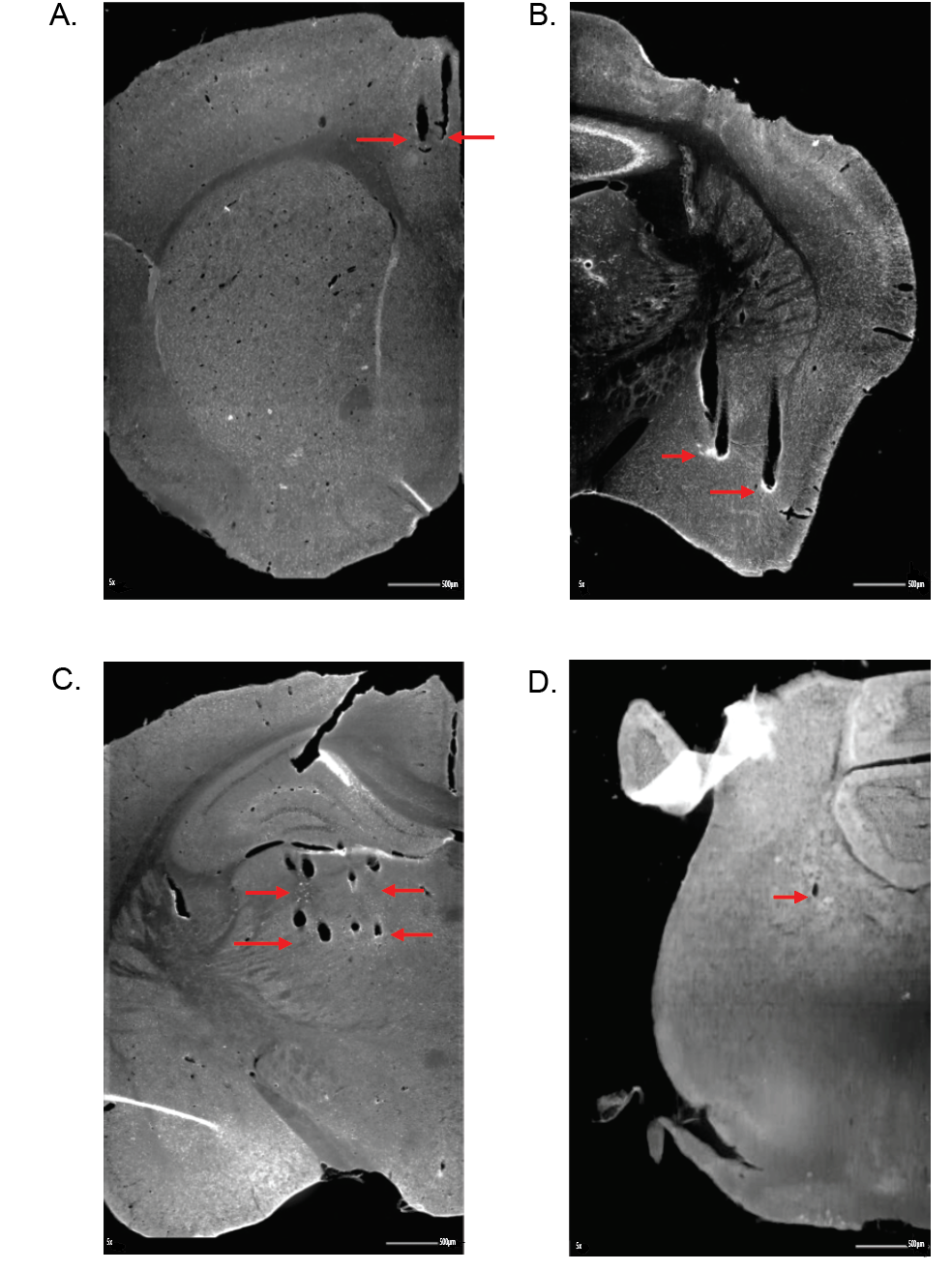

### S1: Histological verification in implanted animals

Electrode wires were verified in the A) ACC, B) Amygdala (BLA/CeA), C) Thalamus (MDthal/Po/VPM), and D) PBN. Red arrows indicate the ends of electrode wire tips. Note, unlike other regions, PBN was targeted at a 15 degree angle so only the very tip is visible, though tracks were traced across multiple slices.

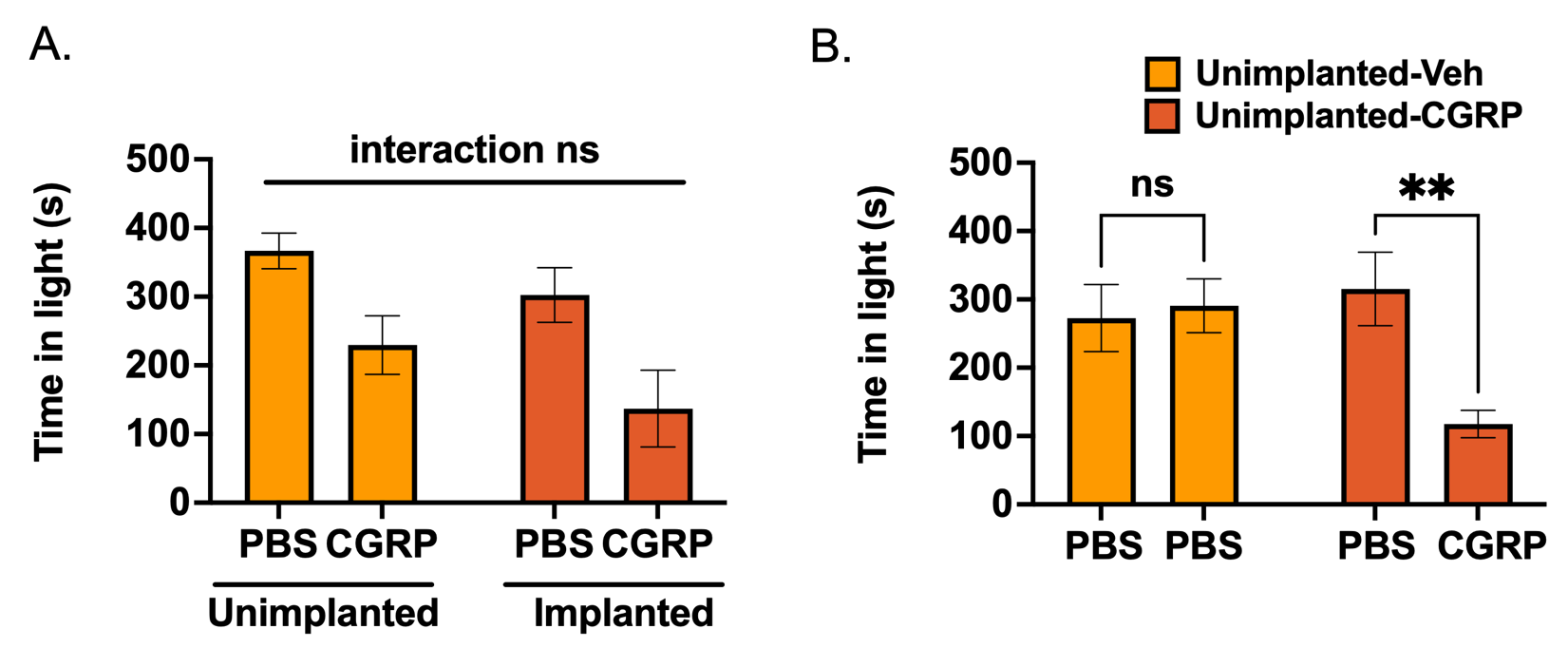

S2: Light Aversion Behavior in implanted and unimplanted animals

Light aversion was assessed using the average amount of total time spent in light (in seconds) over 15 minutes ^1,2^. A repeated measures design was used to compare unimplanted and implanted animals in response to CGRP and performed a control that determined that there is no effect of repeated injection over days using groups that received PBS injections both days and/or CGRP on the treatment day. We used a three-factor mixed-effects model to evaluate the impact of implant, CGRP, and repeated measures. A) Unimplanted (n= 7) and electrode-implanted (n= 7) CD1 mice exhibit light aversive behavior in response to CGRP, and the treatment interaction between the treatment and electrode-implanted condition was not significant. B) A subset of unimplanted mice demonstrate no difference in time spent in light when treated with PBS (n= 5) on the treatment day, while CGRP-treated mice exhibit decreased time spent in light (n= 7). Data are represented as mean ± SEM.

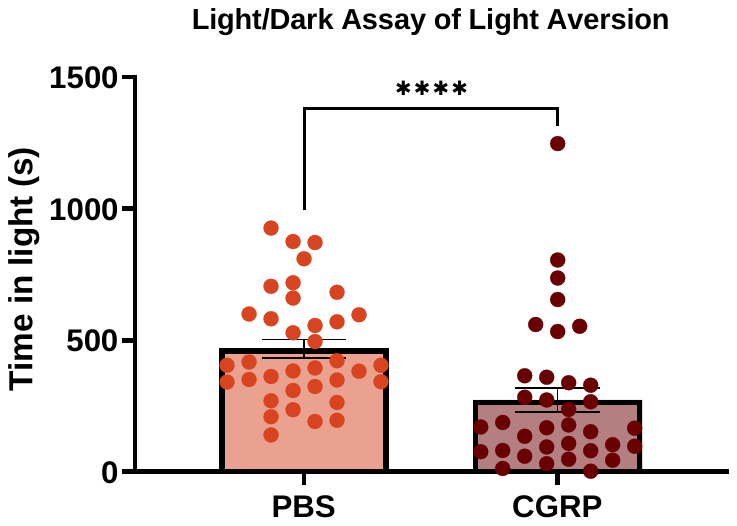

A.

B.

C.

D.

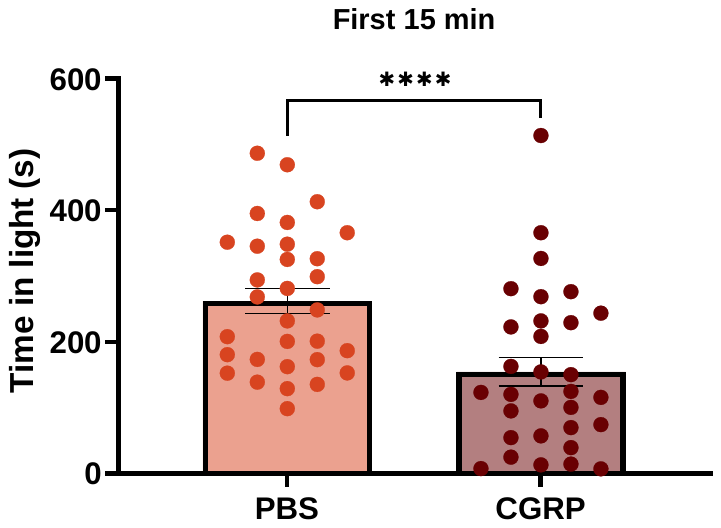

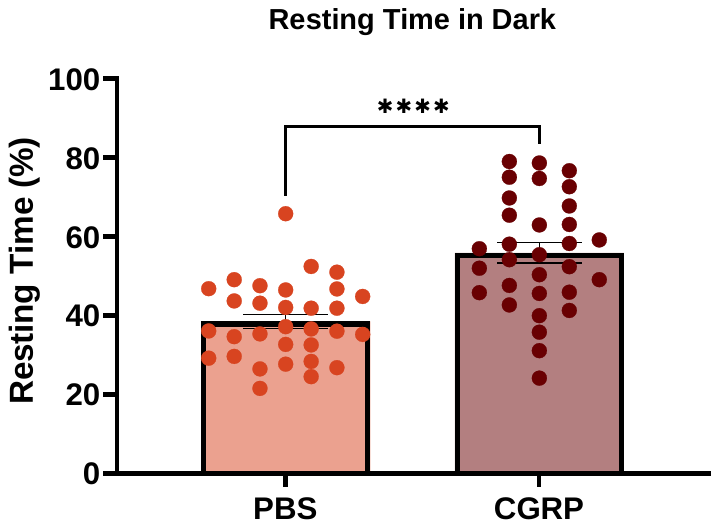

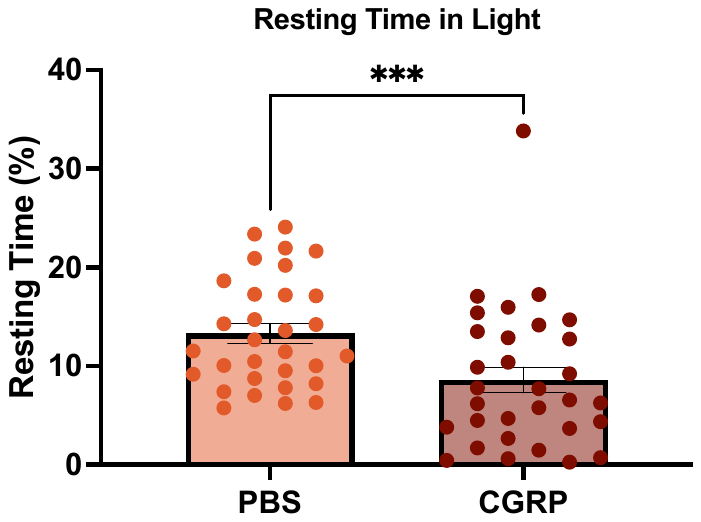

### S3: Light aversion across all implanted animals

In addition to the studies in S2 directly comparing implanted and unimplanted animals, all behavioral measures for each implanted animal are shown here. A and B represent time in light during the total 30-minute duration (A; t(30) = 5.18, p<0.0001) and first 15 minutes (B; t(30) = 5.74, p<0.0001) of the experiment in response to PBS and CGRP. C and D represent the percentage of time spent resting in the dark (C; t(30) = 7.96, p<0.0001) and light (D; t(30) = 4.01, p = 0.0004) over the total amount of time during the first 15 minutes of the light/dark experiment. Two-tailed paired t-tests. ****p<0.0001, **p<0.001. N = 31, Data are represented as mean ± SEM.

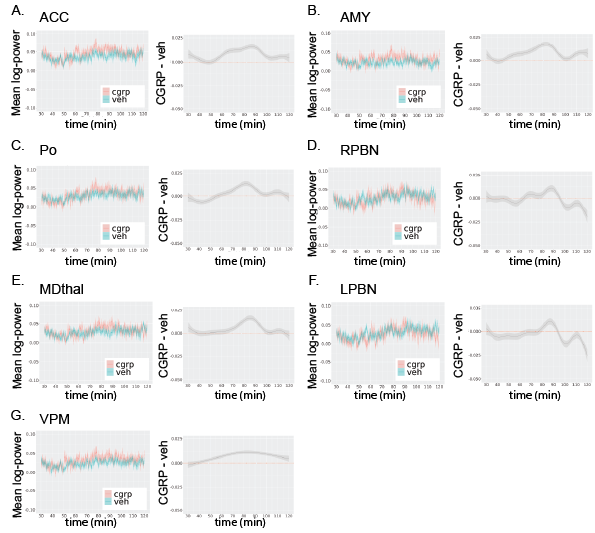

### S4: CGRP Power in 1-3 Hz Frequency Band

Mean log-power is plotted across time following CGRP (pink) and PBS (blue) on left. The difference between the mean log-power across time following CGRP and PBS is shown on the right. FOS analysis revealed significant differences in the: ACC (A; FDR p = 3.74e-16), AMY (B; FDR p=3.74e-16 ), Po (C; FDR p = 3.56e-8), RPBN (D; FDR p = 0.055), MDthal (E; FDR p = 3.74e-16), LPBN (F; FDR p = 8.88e-9), and VPM (G; FDR p = 3.74e-16). Data are represented as 30-second rolling means ± SEM.

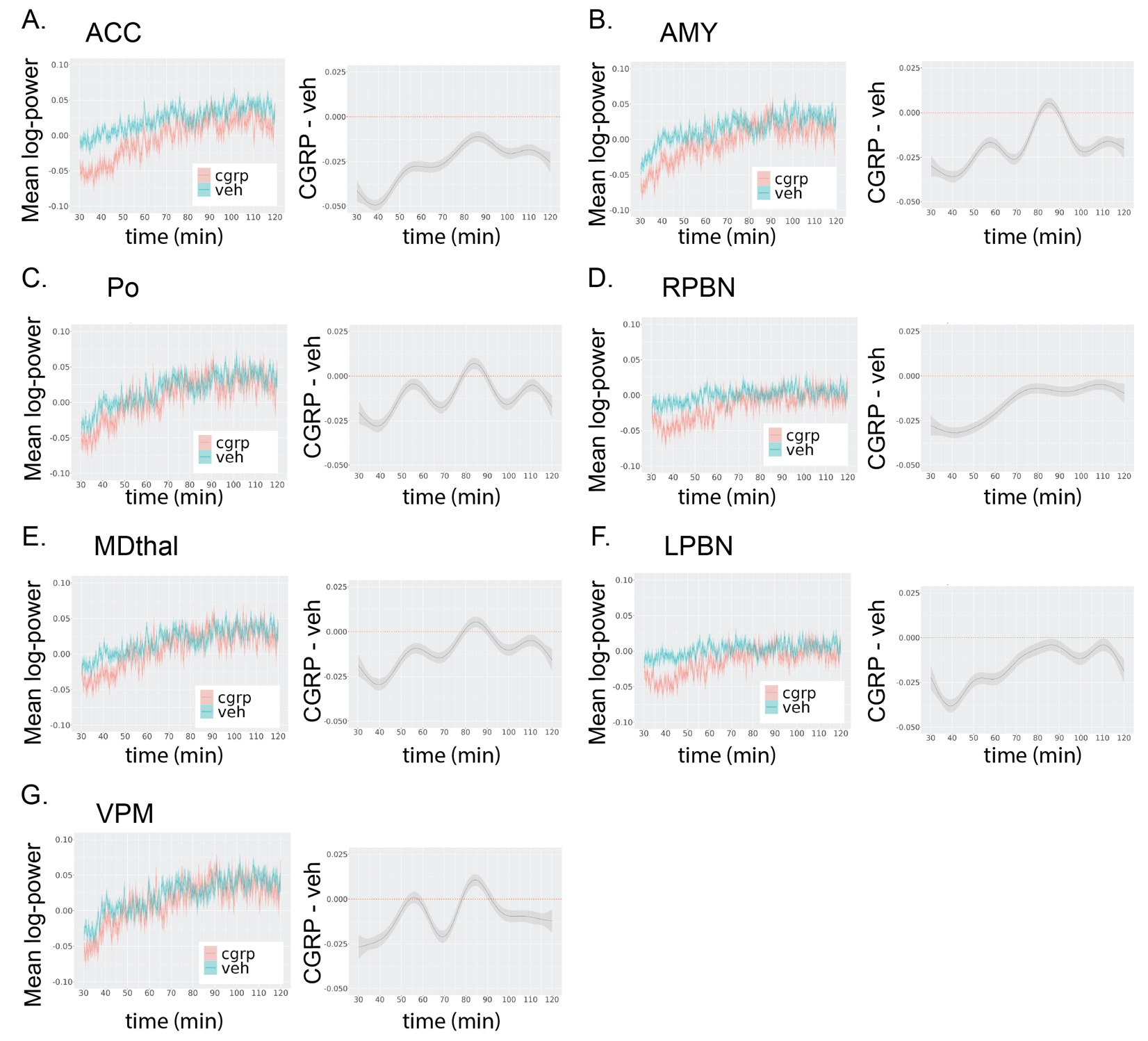

### S5: CGRP Power in 8-13 Hz Frequency Band

Mean log-power is plotted across time following CGRP (pink) and PBS (blue) on left. The difference between the mean log-power across time following CGRP and PBS is shown on the right. FOS analysis revealed significant differences in the: ACC (A; FDR p = 3.74e-16), AMY (B; FDR p =3.74e-16), Po (C; FDR p = 3.74e-16), RPBN (D; FDR p = 3.74e-16), MDthal (E; FDR p = 3.74e-16), LPBN (F; FDR p = 3.74e-16), and VPM (G; FDR p = 3.74e-16). Data are represented as 30-second rolling means ± SEM.

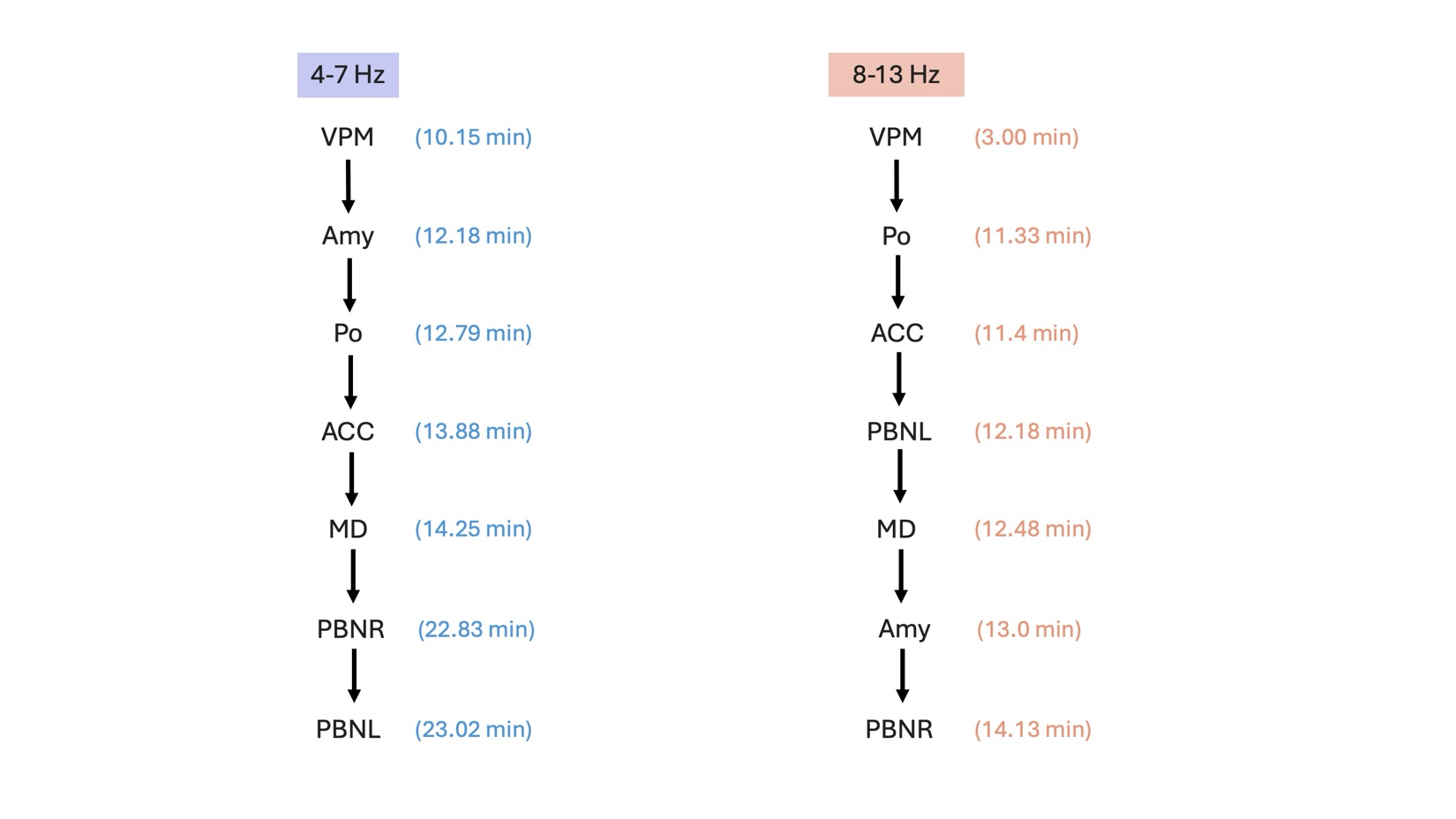

### S6 Timing order of power peak magnitude differences between CGRP and vehicle

Time (in minutes) of peak magnitude difference follow injection between CGRP and vehicle shown in parentheses. VPM power peaks first at both 4-7Hz and 8-13Hz.

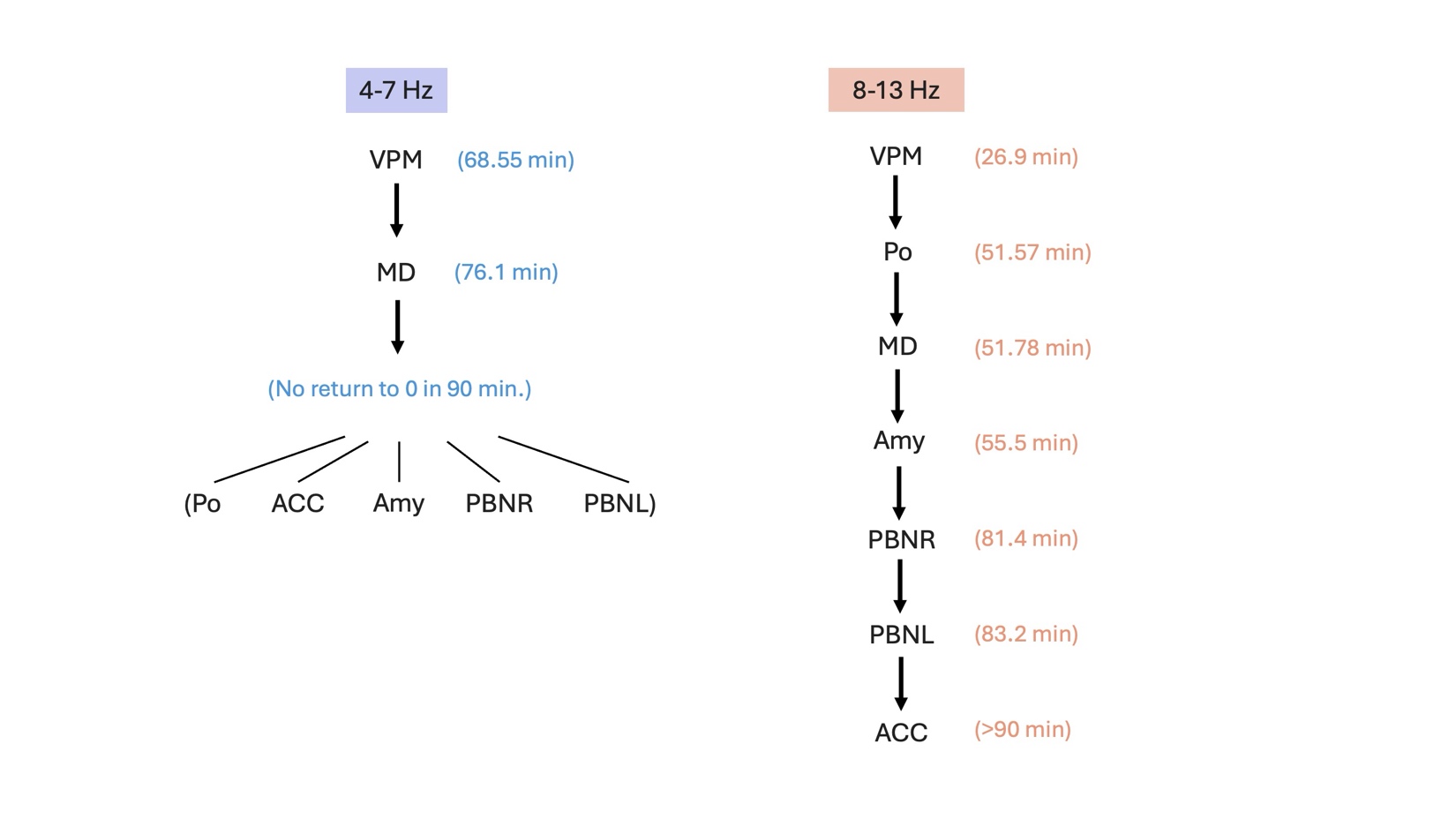

### S7 Timing order of power return to zero differences between CGRP and vehicle

Time (in minutes) following injection until return to zero difference between CGRP and vehicle shown in parentheses. VPM power is the first to return to zero at both 4-7Hz and 8-13Hz. At 4-7Hz, five regions do not return to zero within the 90 minutes post-injection measured.

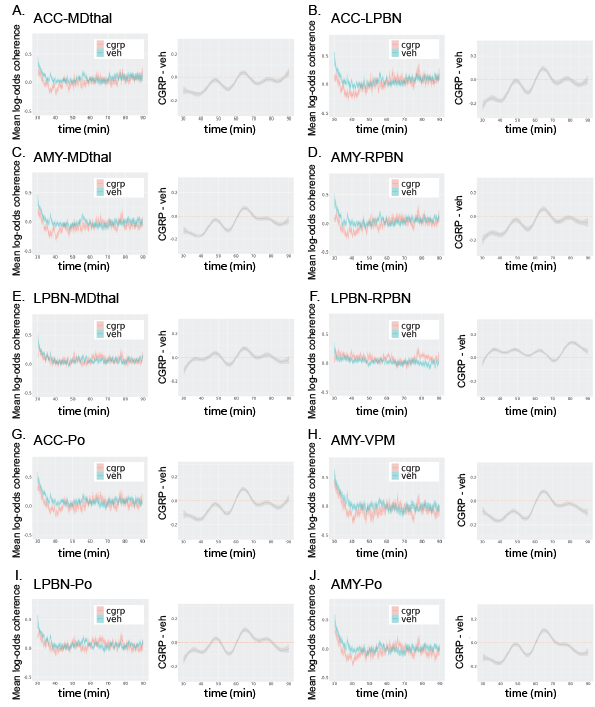

### S8: CGRP Coherence in 4-7 Hz Frequency Band

Mean log-odds coherence is plotted across time following CGRP (pink) and PBS (blue) on left. The difference between the mean log-odds coherence following CGRP and PBS is shown on the right. FOS analysis revealed significant differences in the: ACC-MDthal (A; FDR p = 3.74e-16), ACC-LPBN (B; FDR p = 3.74e-16), AMY-MDthal (C; FDR p = 3.74e-16), AMY-RPBN (D; FDR p = 3.74e-16), LPBN-RPBN (F; FDR p = 3.74e-16), ACC-Po (G; FDR p = 3.74e-16), AMY-VPM (H; FDR p = 3.74e-16), LPBN-Po (I; FDR p = 2.05e-7), and AMY-Po (J; FDR p = 3.74e-16). There was no significant difference in the LPBN-MDthal (E; FDR p = 0.803). Data are represented as 30-second rolling means ± SEM.

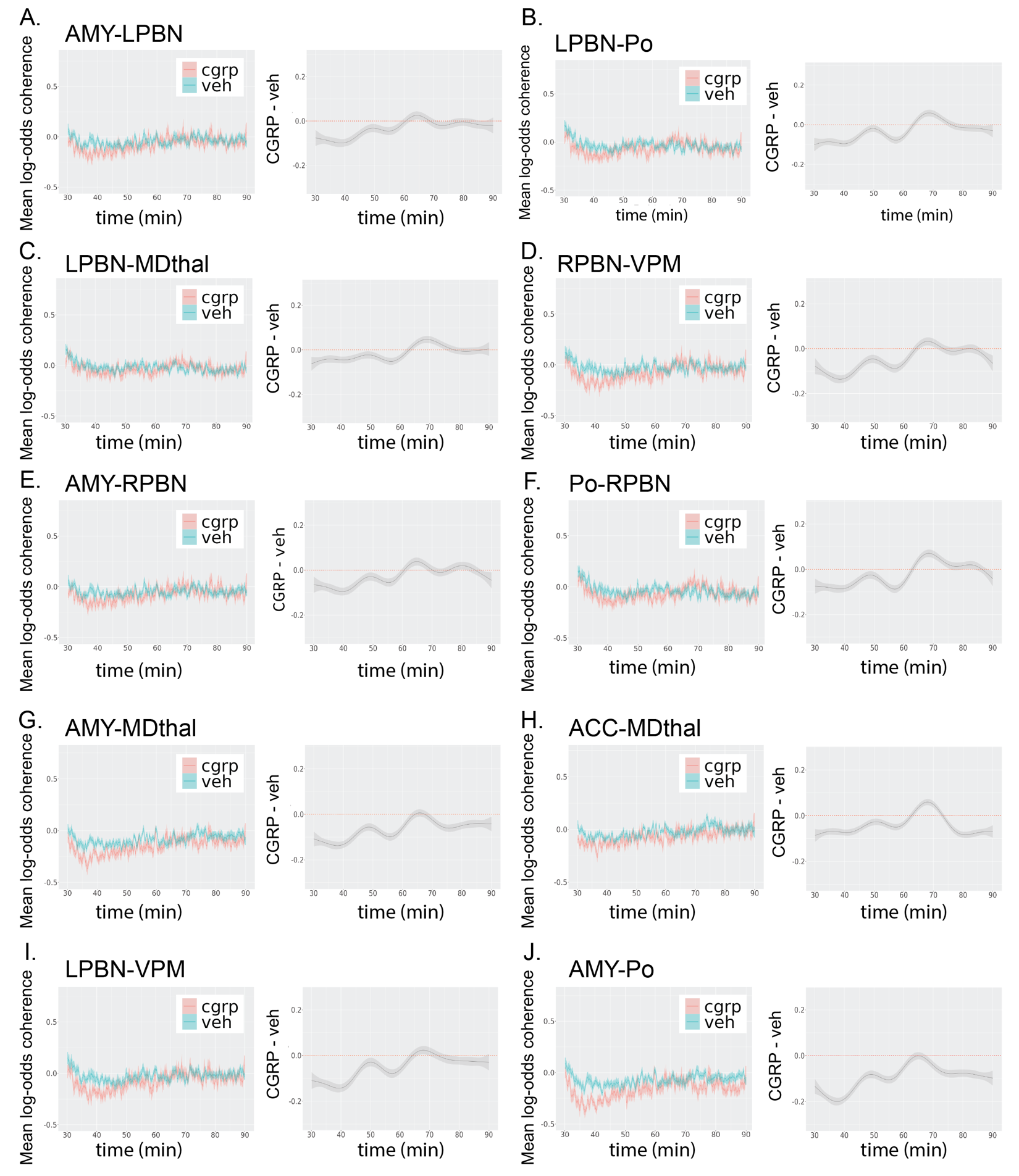

### S9: CGRP Coherence in 8-13 Hz Frequency Band

Mean log-odds coherence is plotted across time following CGRP (pink) and PBS (blue) on left. The difference between the mean log-odds coherence following CGRP and PBS is shown on the right. FOS analysis revealed significant differences in the: AMY-LPBN (A; FDR p = 3.74e-16), LPBN-Po (B; FDR p = 3.74e-16), LPBN-MDthal (C; FDR p = 1.22e-7), RPBN-VPM (D; FDR p = 3.74e-16), AMY-RPBN (E; FDR p = 3.74e-16) , Po-RPBN (F; FDR p = 9.59e-16), AMY-MDthal (G; FDR p = 3.74e-16), ACC-MDthal (H; FDR p = 3.74e-16), LPBN-VPM (I; FDR p = 3.74e-16), and AMY-Po (J; FDR p = 3.74e-16). Not shown and significant: ACC-Po (FDR p = 3.74e-16), ACC-VPM (FDR p = 3.74e-16), AMY-VPM (FDR p = 3.74e-16), LPBN-Po (FDR p = 3.74e-16), MDthal-RPBN (FDR p = 0.0219), MDthal-VPM (FDR p = 3.74e-16), MDthal-Po (H; FDR p = 3.74e-16), and Po-VPM (FDR p = 3.74e-16). Not shown and not significant: ACC-LPBN (FDR p = 0.237), ACC-RPBN (FDR p = 0.179), and LPBN-RPBN (FDR p = 0.137). Data are represented as 30-second rolling means ± SEM.

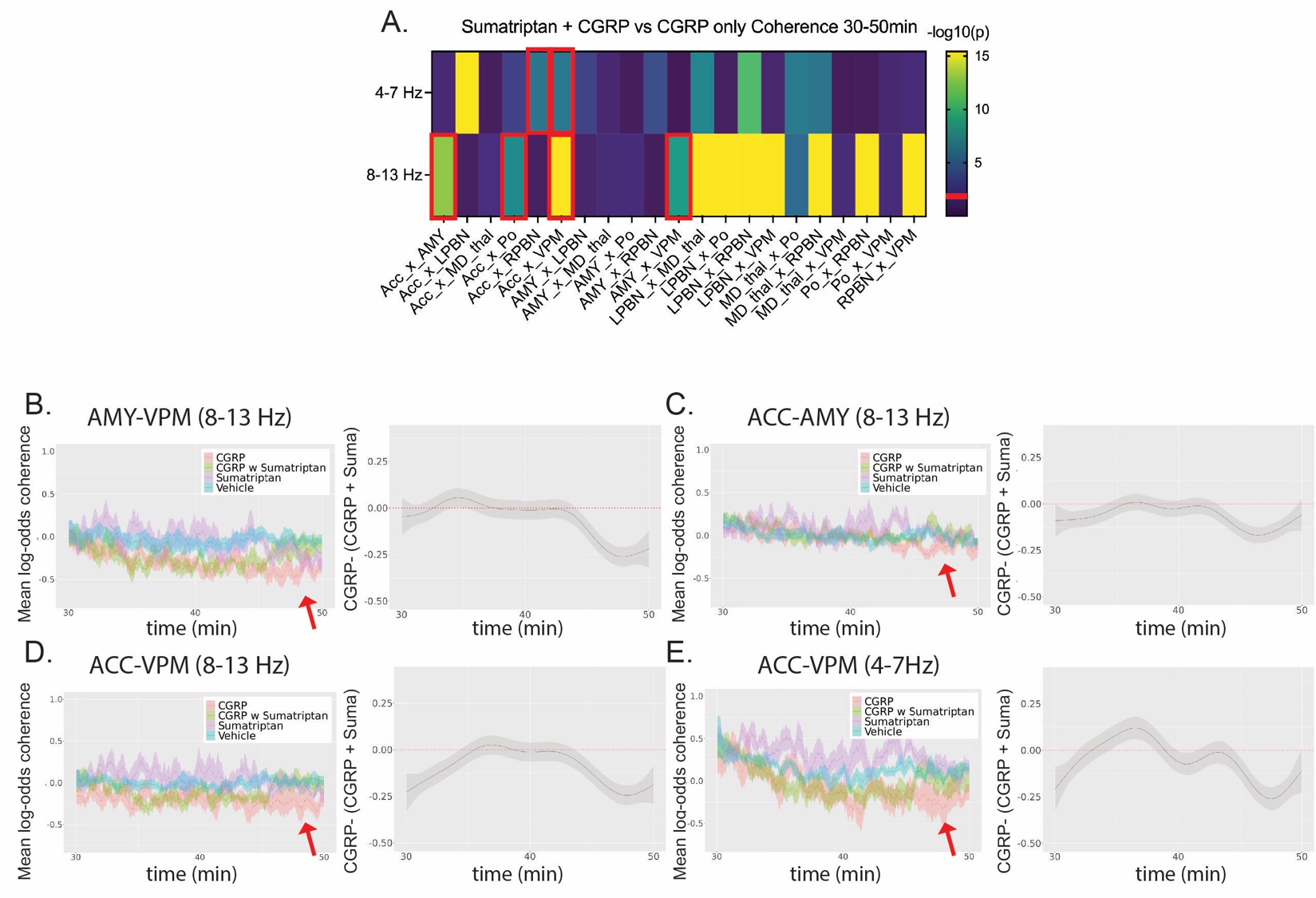

### S10: Partial rescue of CGRP-induced brain network coherence by Sumatriptan

A) Heat map showing -log10 p-values summarizing the effect of sumatriptan+CGRP on LFP coherence. Red boxes indicate significant differences in which reversal of the CGRP effect occurred. B-E) The mean log-odds coherence are plotted for each condition across time (left) and difference in mean log-odds coherence between sumatriptan+CGRP and CGRP conditions (right) for region pairings. Coherence data are represented as 60-second rolling means ± SEM. Mean logpower and mean log-odds coherence (C-H) is plotted across time during the following conditions: CGRP alone (red), CGRP with sumatriptan (green), sumatriptan (purple), and vehicle (blue).

With regard to the three brain region pairings where CGRP was found to most disrupt coherence, we observed significant diminution of the CGRP response by sumatriptan in coherence between the AMY-VPM 8-13Hz (FDR p=2.57e-9) but not 4-7 Hz (FDR p=0.69, Figure S11), ACC-LPBN 4-7 Hz (FDR p=2.92e-9, Figure S11), but not 8-13 Hz (FDR p=0.7) and AMY-LPBN 4-7Hz (FDR p= 8.92e-4, Figure S11) but not 8-13Hz (FDR p=0.1, Supplemental Figure S10).

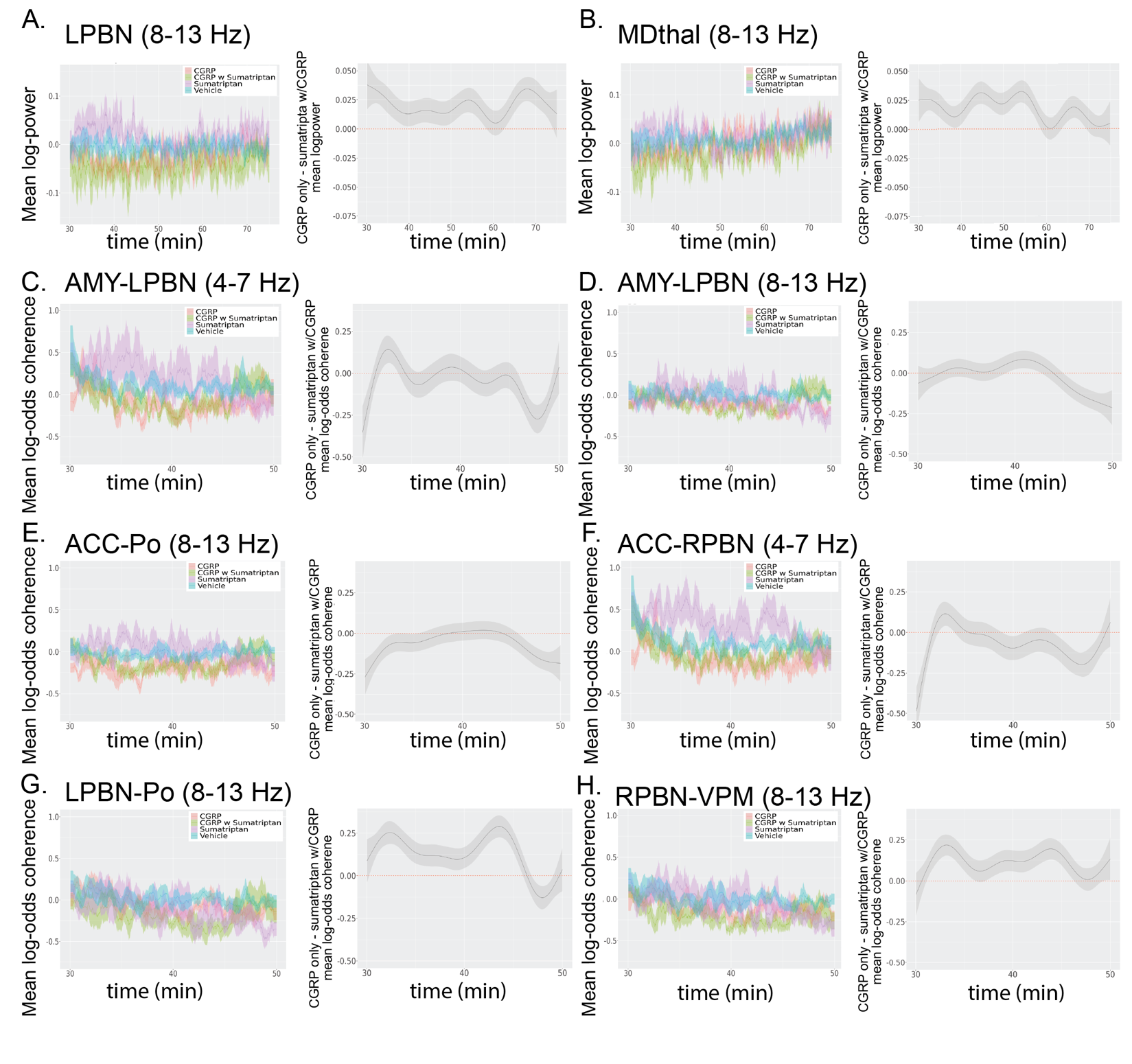

### S11: Additional Sumatriptan power and coherence responses

A-B) Mean logpower is plotted across time during the following conditions: CGRP alone (red), CGRP with sumatriptan (green), sumatriptan (purple), and vehicle (blue). Sumatriptan+CGRP coadministration induced similar changes in mean logpower as seen in CGRP-only administration in the LPBN (A; FDR p = 3.74e-16), and MDthal (B; FDR p = 3.74e-16).

C-H) Mean log-odds coherence is plotted for each brain region pairing. Sumatriptan+CGRP reverses coherence changes observed in the AMY-LPBN in the 4-7 Hz frequency band (C; FDR p = 8.29e-4), but not in the 8-13 Hz frequency band for this brain region pairing (D; FDR p = 0.105). Sumatriptan+CGRP coadministration reverses coherence changes observed in the ACC-Po in the 8-13 Hz frequency band (E; FDR p = 4.68e-8) and in the ACC-RPBN in the 4-7 Hz frequency band (F; FDR p = 1e-6). However, sumatriptan+CGRP induced coherence changes in the same direction as CGRP for parabrachial-thalamic pairings (G-H; FDR p= 3.74e-16).

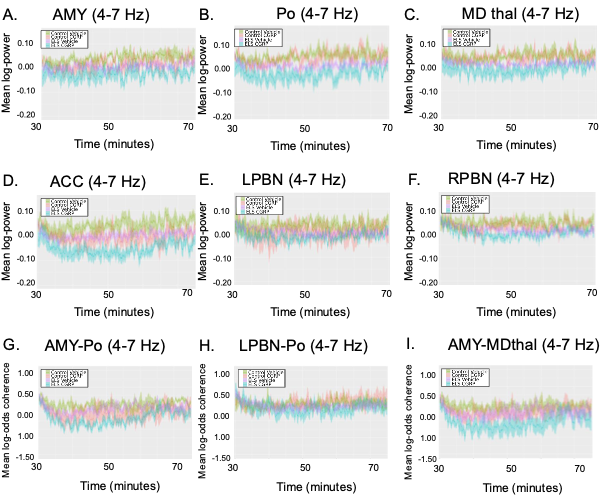

### S12: Early life stress power and coherence in response to CGRP in 4-7 Hz band

A-B) Mean logpower and mean log-odds coherence (C-D) is plotted across time during the following conditions: CGRP with control animals (red), vehicle with control animals (green), ELS CGRP (blue), and ELS vehicle (purple). There were significant interactions between ELS and CGRP with regard to power in the AMY (A; FDR p = 3.88e-3), Po (B; FDR p = 3.74e-16), MD Thal(C; FDR p=3.74e-16), ACC (D; FDR p=1.05e-4), LPBN (E; FDR p= 3.71e-06), but not the RPBN(F; FDR p= 0.49). Significant differences in ELS coherence were observed in the AMY-Po (G; FDR p = 6.03e-4) and LPBN-Po (H; FDR p = 1.93e-15), AMY-MDthal (I; FDR p=3.74e-16).

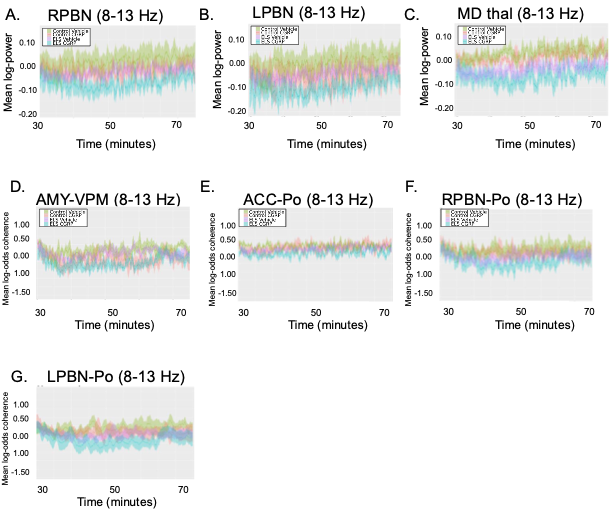

### S13: Early life stress power and coherence in response to CGRP in 8-13 Hz band

A-C) Mean logpower and D-F) mean log-odds coherence is plotted across time during the following conditions: CGRP with control animals (red), vehicle with control animals (green), ELS CGRP (blue), and ELS vehicle (purple). There were significant differences in ELS power in the RPBN (A; FDR p=.002) and MDthal (C; FDR p = 1.13e-16), but not the LPBN (B; FDR p= 0.19). (D-F) Significant differences in ELS coherence in response to CGRP were observed in the ACC-Po (E; FDR p = 1.31e-8), RPBN-Po (F; FDR p =2.69e-14), and LPBN-Po (G, FDR p= 5.65e-5). There were no significant differences found in LPBN power (B; FDR p = 0.19) or AMY-VPM coherence (D; FDR p = 0.83).

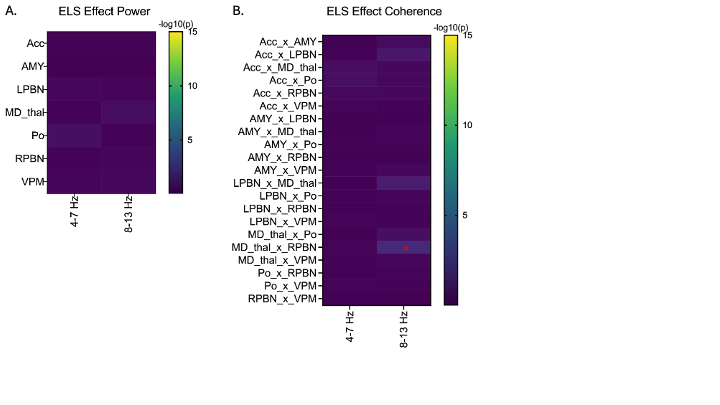

### S14: Heat map of ELS effects

Heat maps showing the effect of ELS when considered in the absence of interaction with CGRP on A) LFP power and B) coherence. ELS largely does not induce significantly different changes in LFP power or coherence when evaluated, MDthalxRPBN FDR-corrected *p<.05, MDthalxLPBN FDR-corrected p=.07.

A.

B.

C.

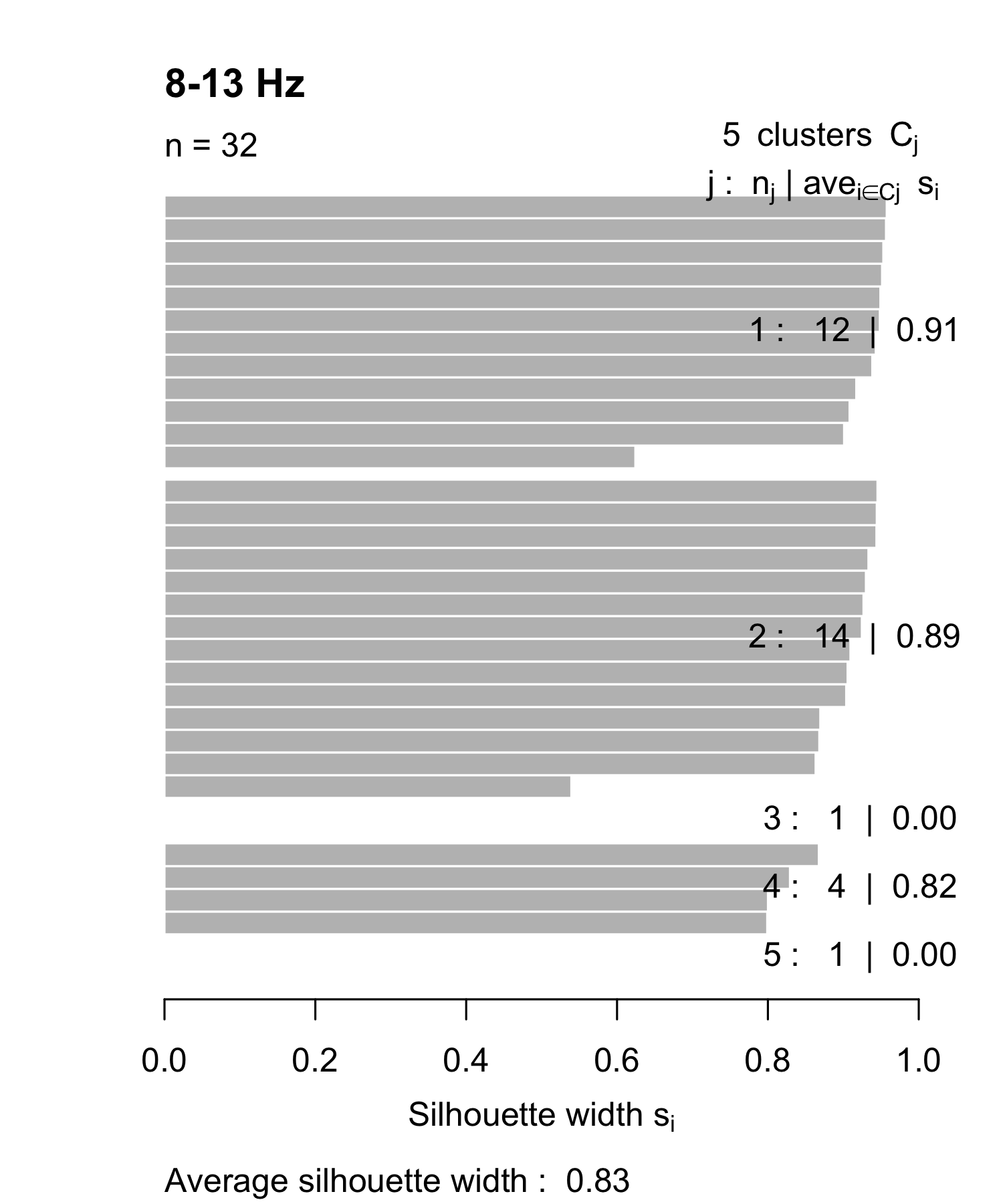

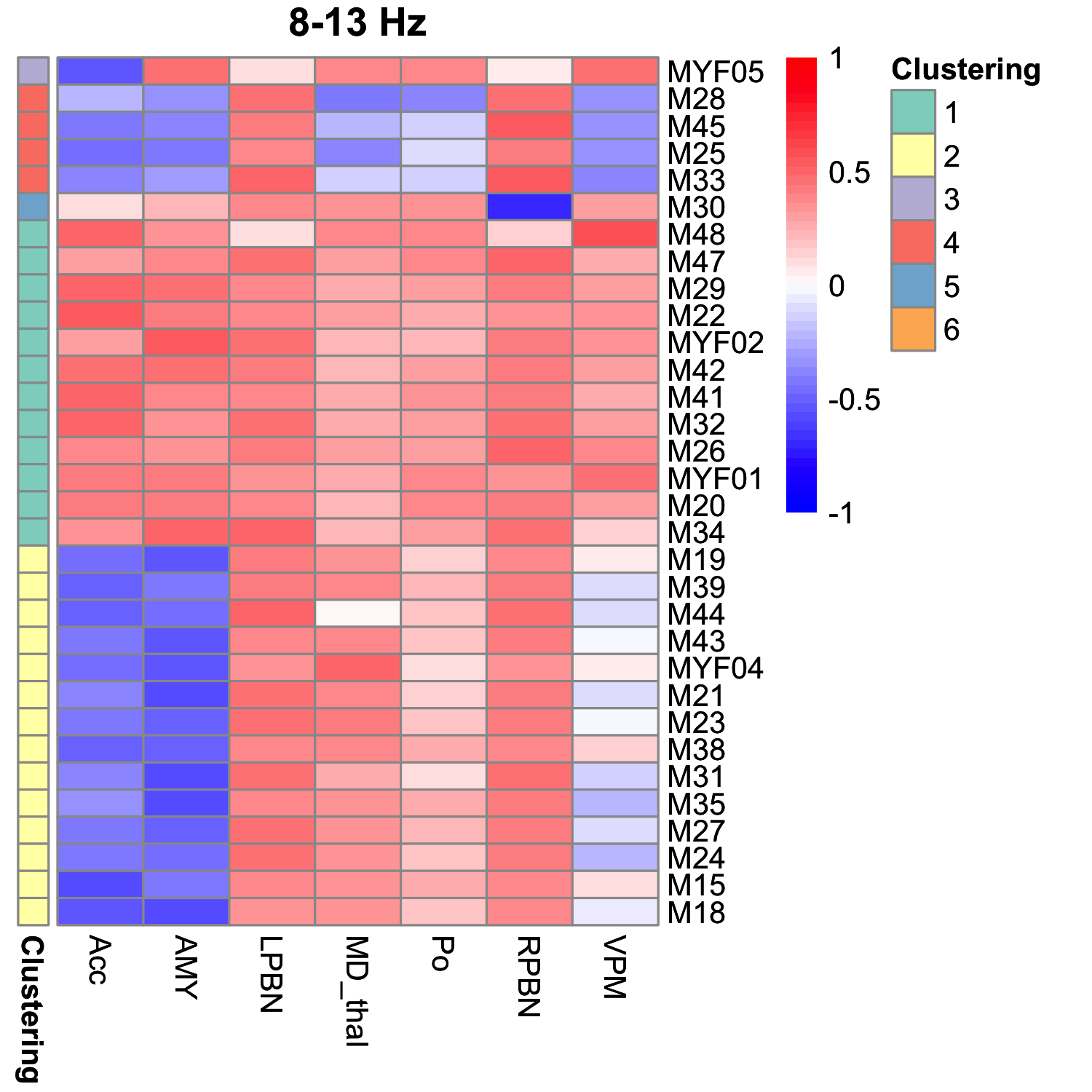

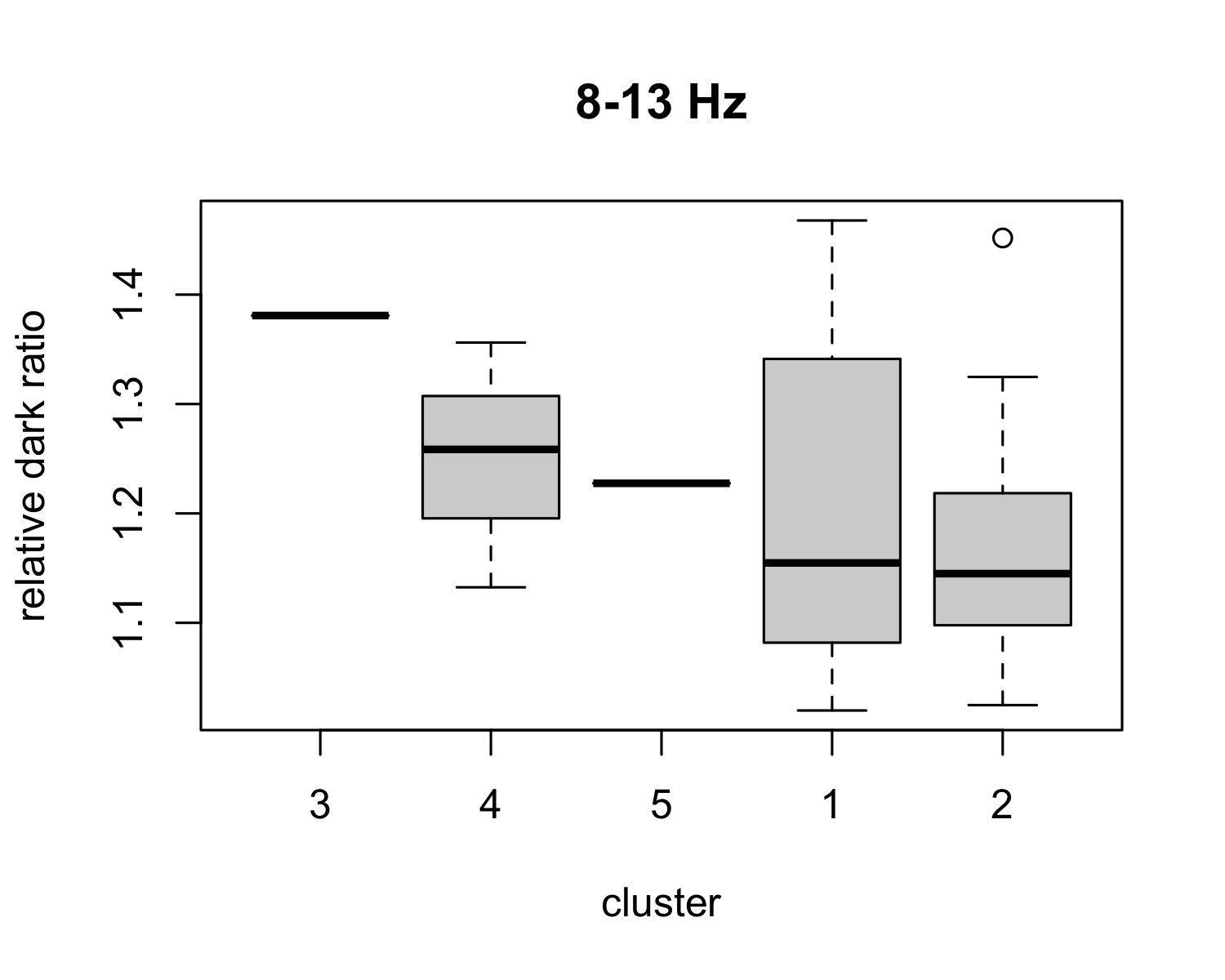

### S15: Changepoint Analysis in 8-13 Hz Frequency Band

A) 8-13Hz k-medoids clustering of initial changepoint projection features following injection with CGRP, silhouette width on the x axis, and each mouse on the y axis, the counts and average silhouette width in each cluster listed on the right, and the overall average silhouette width on the bottom. B) 8-13 Hz projection features for each mouse by cluster. Brain regions are represented across the bottom with each mouse representing a different row, ordered by the cluster average of relative dark ratio. The legend shows the color of values of each cluster in the heatmap. C) 8-13 Hz relationship between brain network feature clustering and behavior on the light/dark test, represented by the relative dark ratio= Treatment time in dark/ Vehicle time in dark.

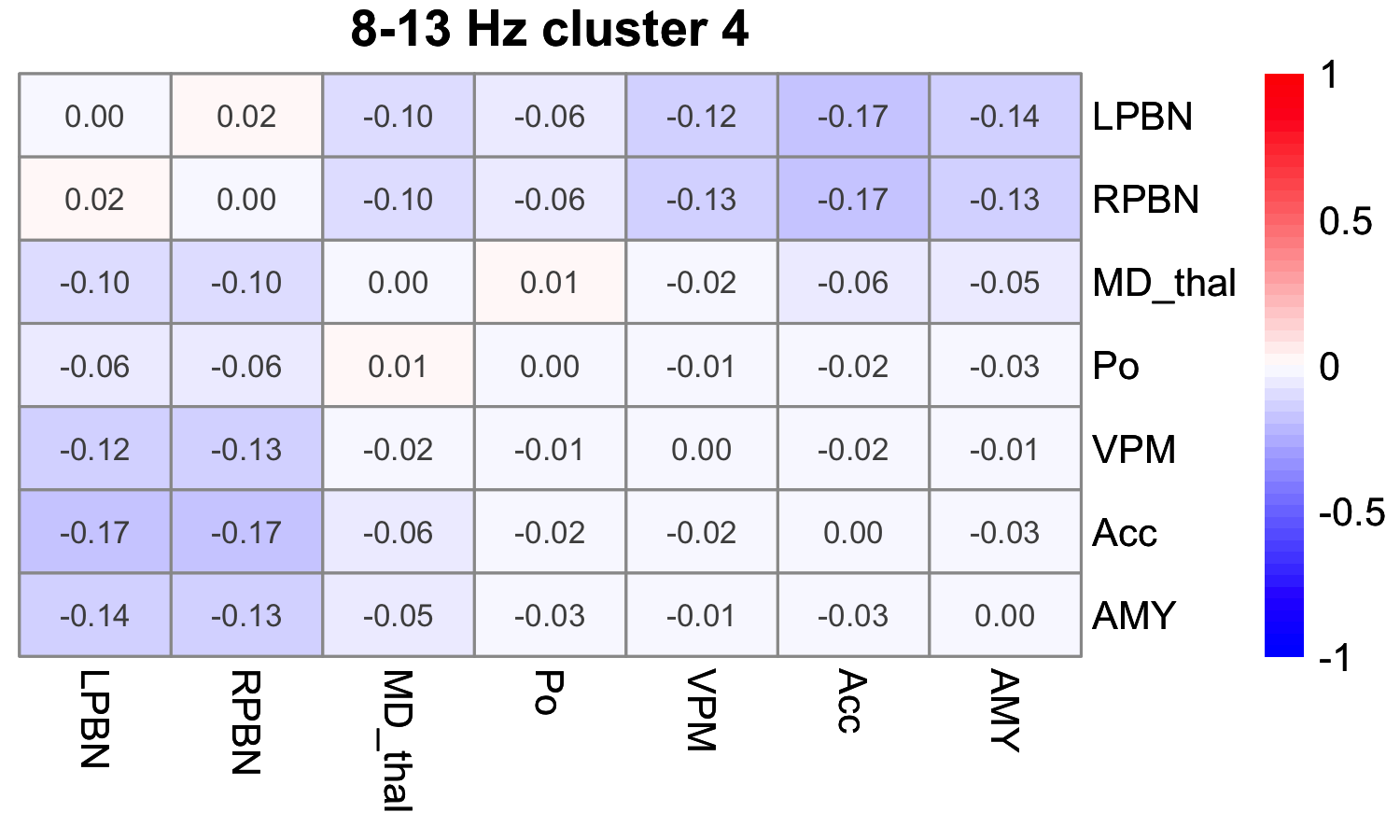

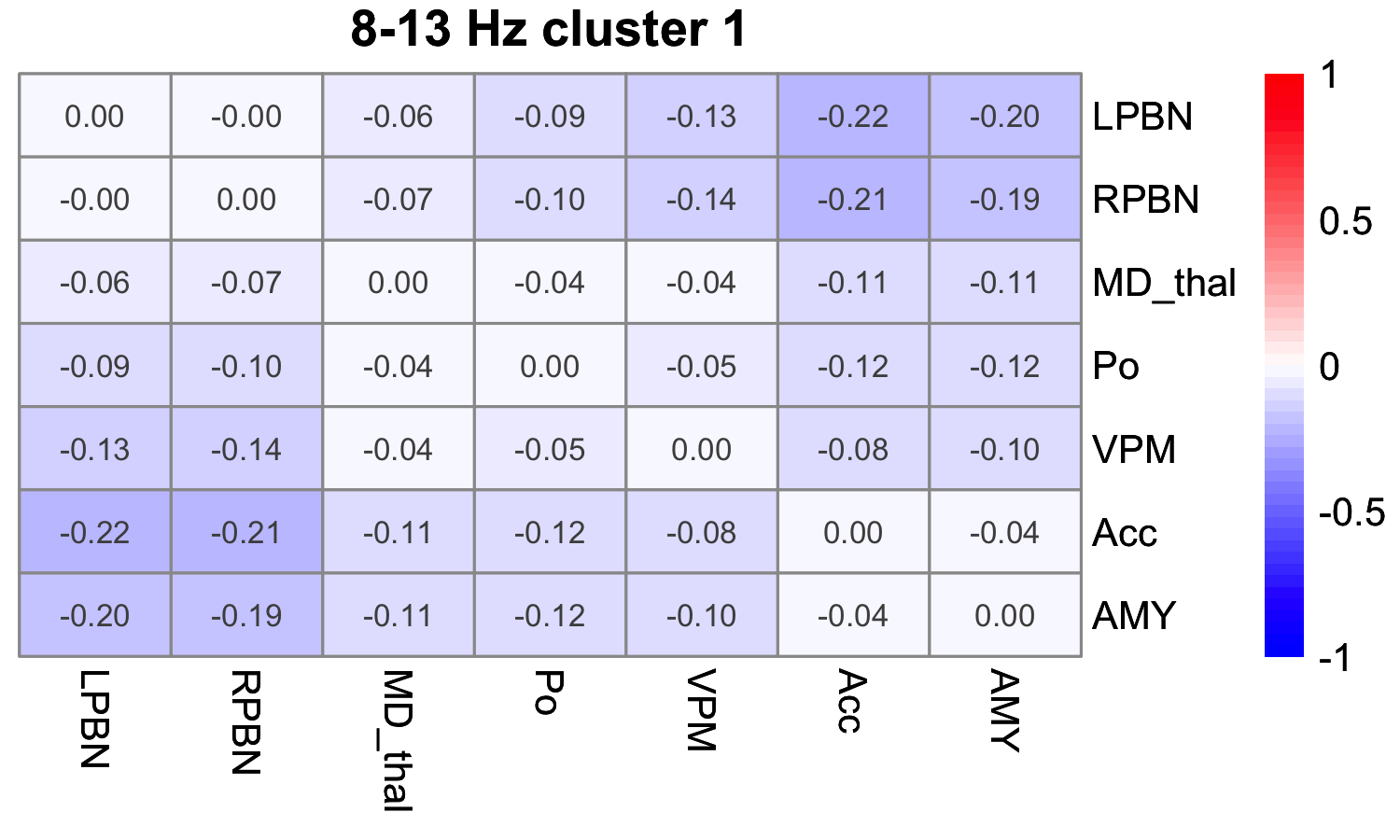

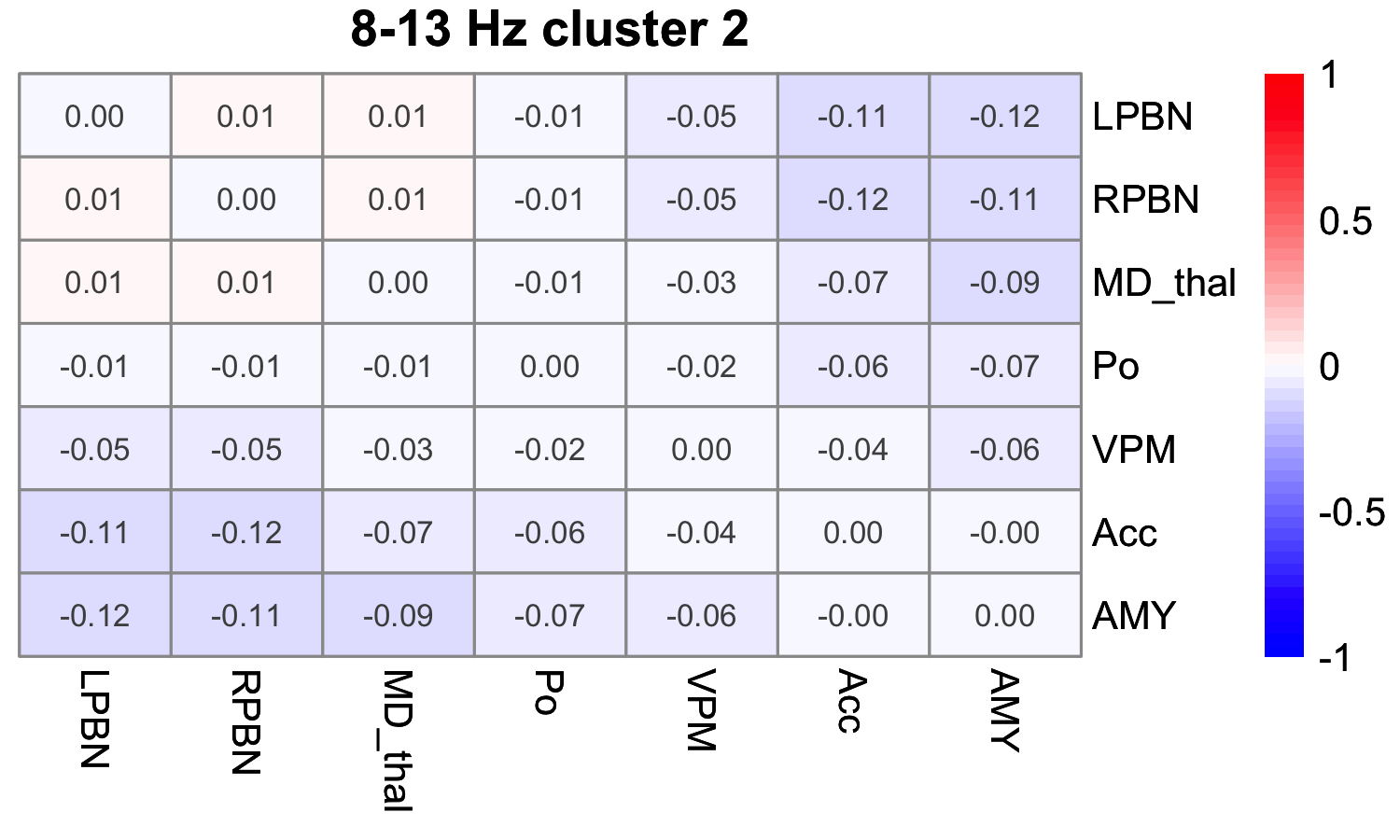

### S16: Coherency Matrices of Changepoint in 8-13 Hz Frequency Band

The cluster mean of difference of coherency matrix is shown for 8-13 Hz over the first change point (i.e. the change in coherency matrices from the segment preceding changepoint to the segment after changepoint) after injection (in clusters with at least 3 mice and ordered by the average relative dark ratio).

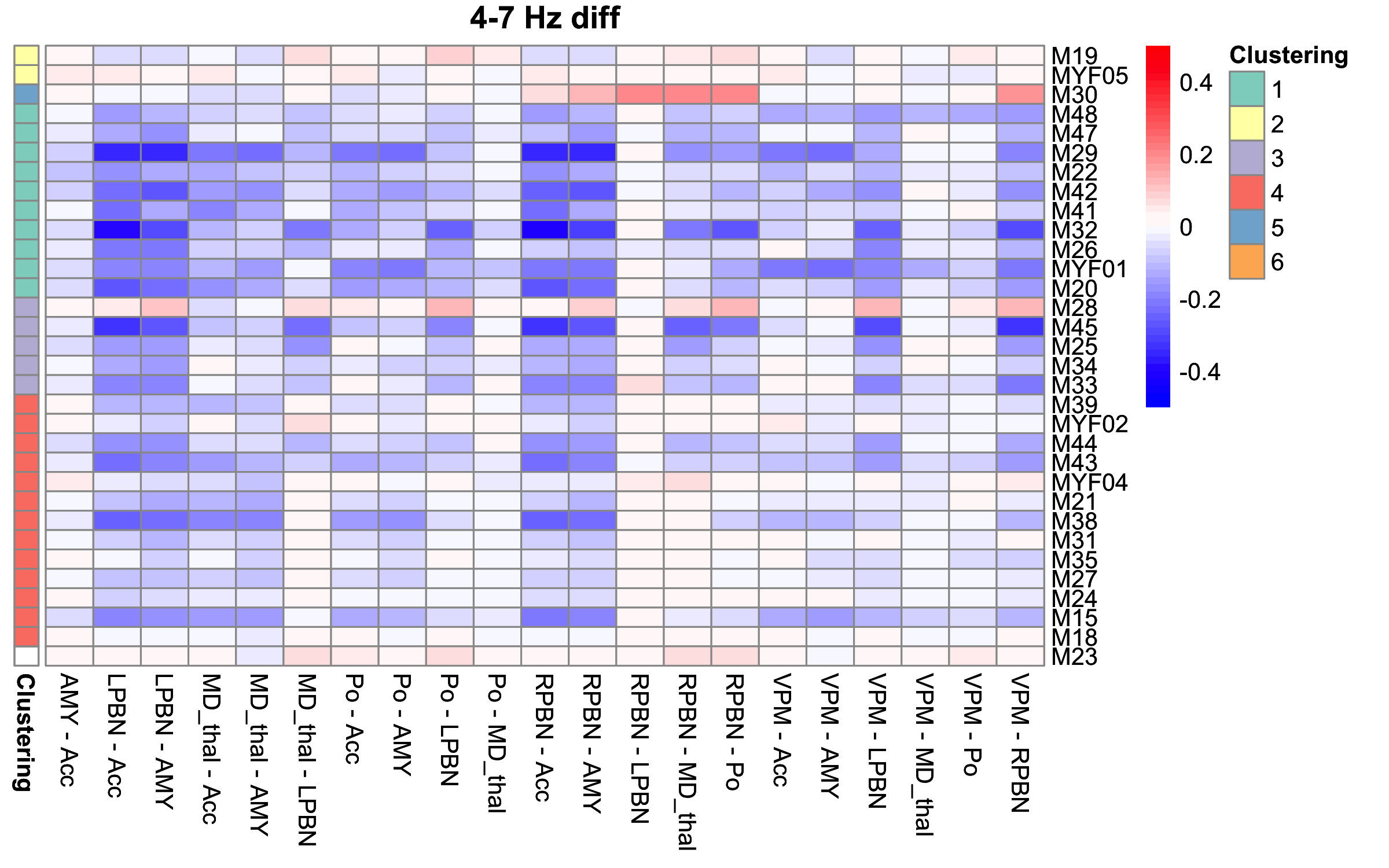

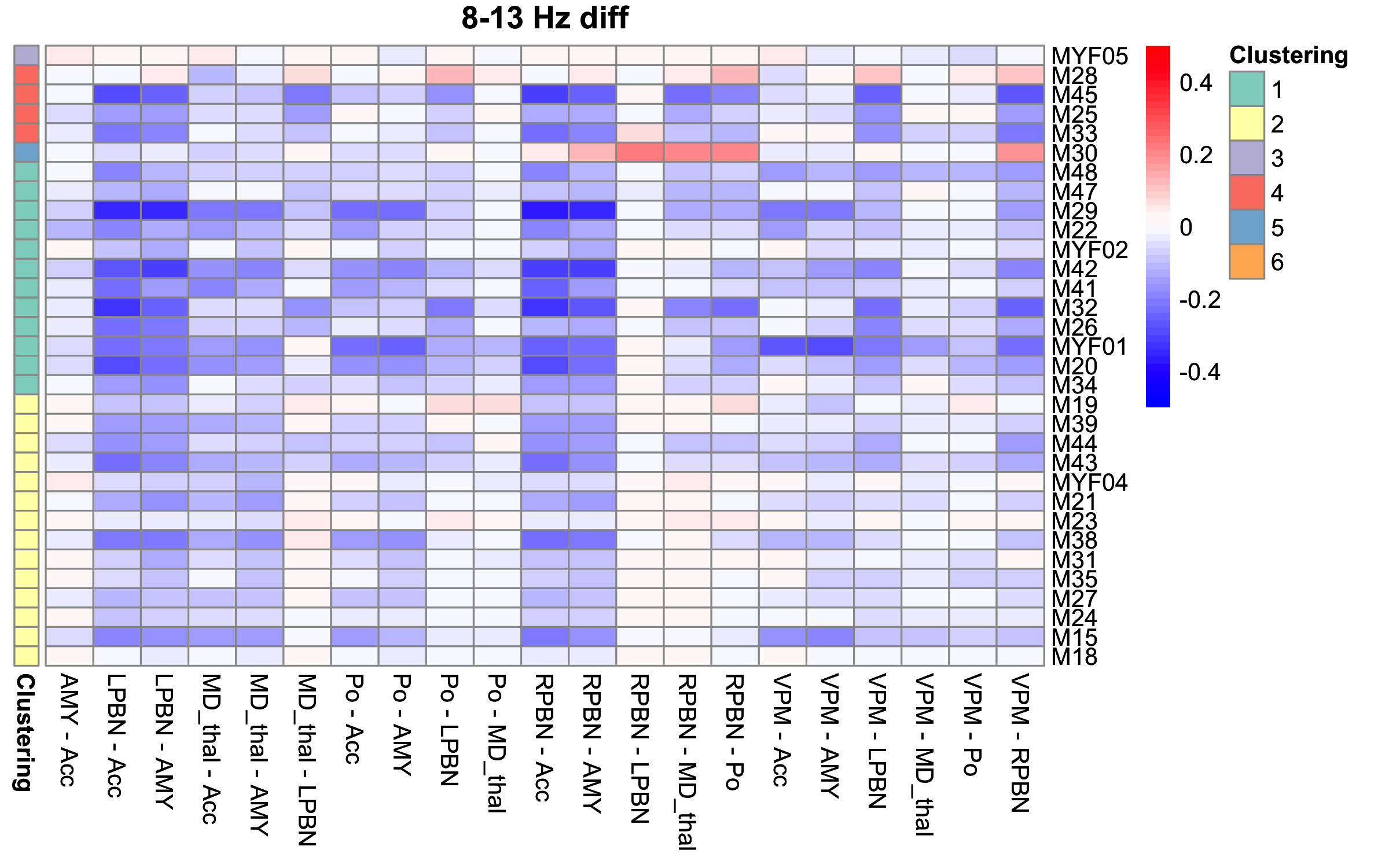

### S17: Coherency Matrices of Changepoint

The difference of coherency matrix on 4-7 Hz (Top) and 8-13 Hz (Bottom) over the first change point after injection for each mouse (i.e. the change in coherency matrices from the segment preceding changepoint to the segment after changepoint).

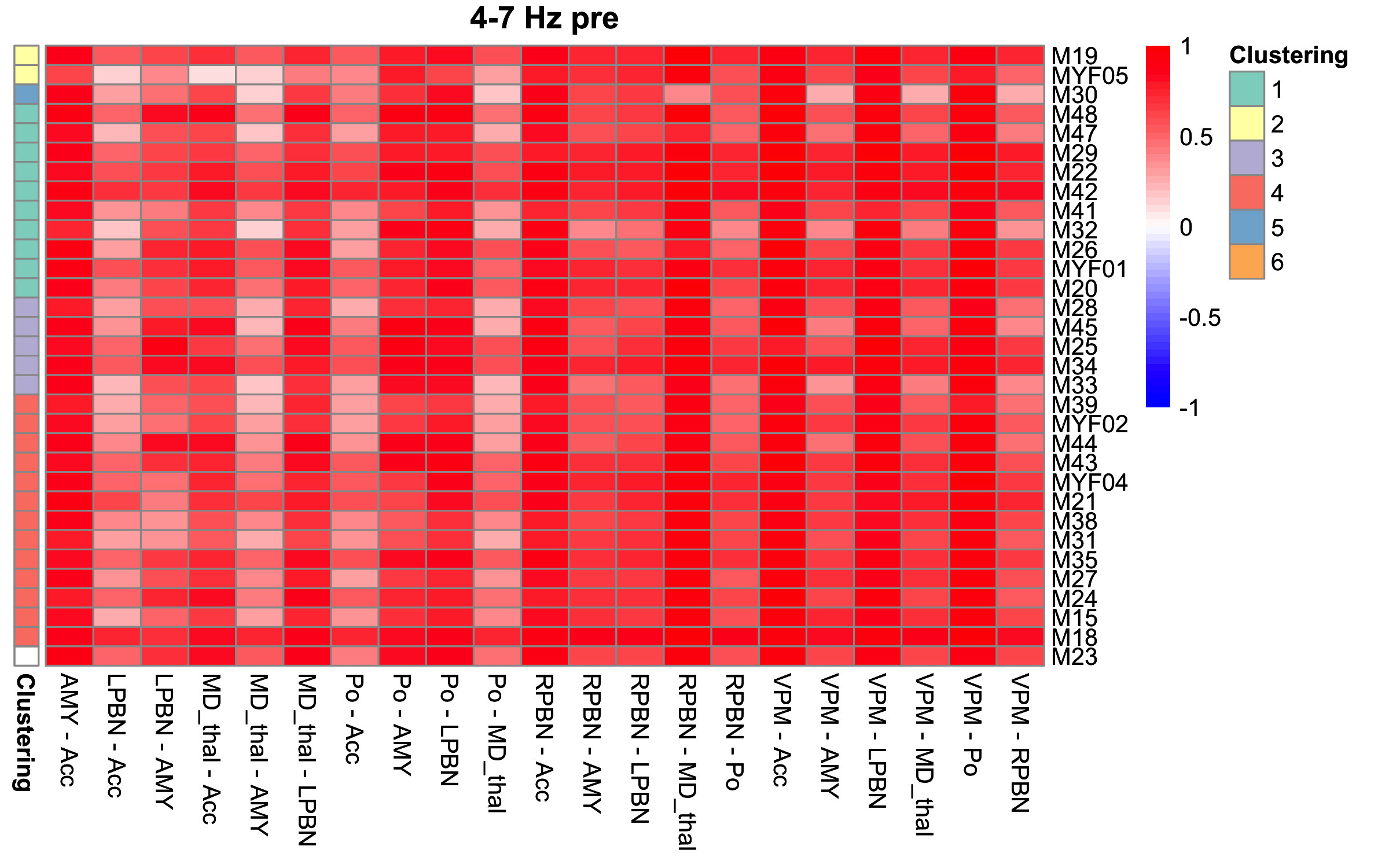

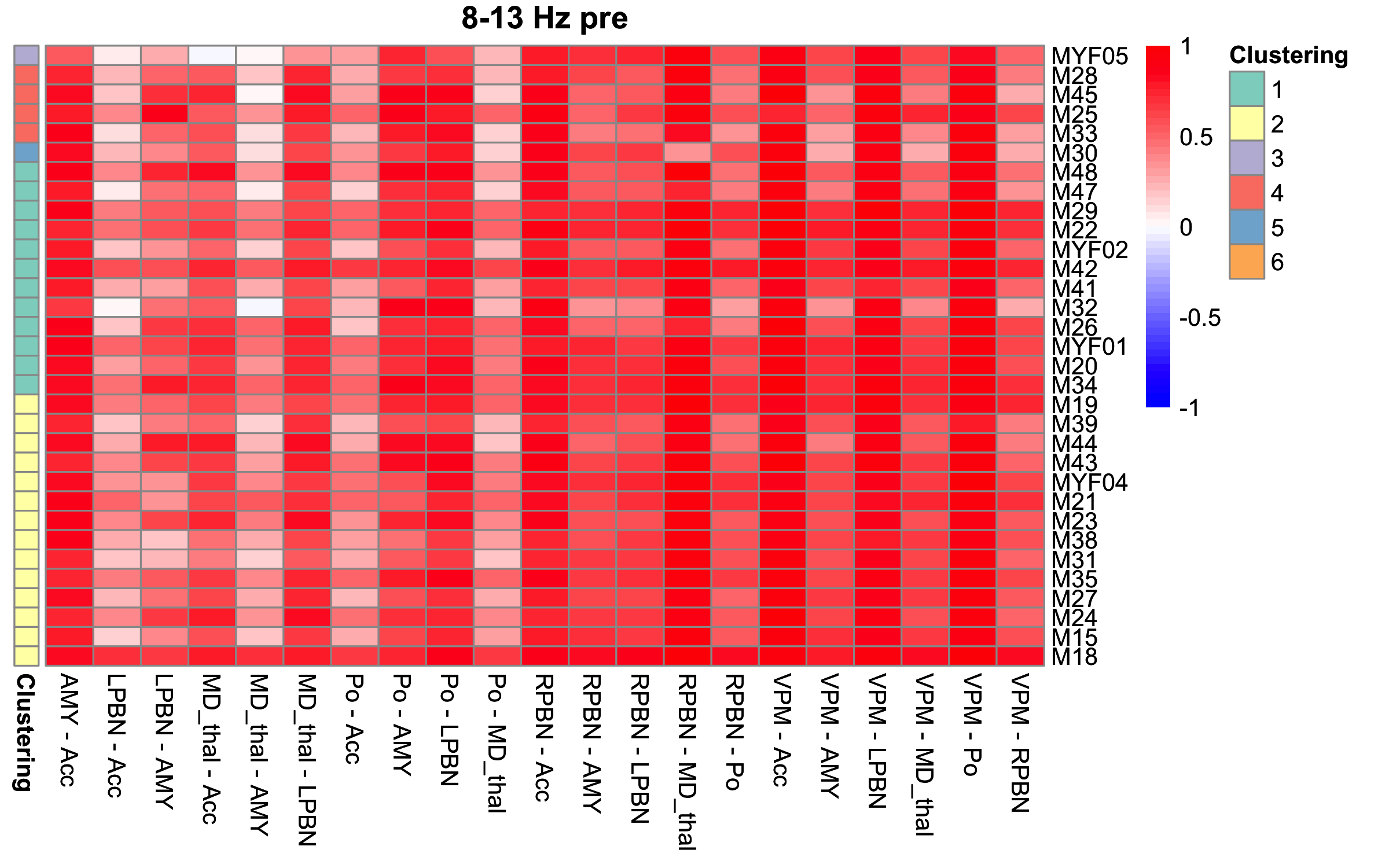

### S18: Coherency Matrices Prior to Changepoint

The coherency matrices at 4-7 Hz (top) and 8-13 Hz (bottom) before the first change point after injection for each mouse.

1.	Mason BN, Kaiser EA, Kuburas A, et al. Induction of Migraine-Like Photophobic Behavior in Mice by Both Peripheral and Central CGRP Mechanisms. *The Journal of neuroscience : the official journal of the Society for Neuroscience*. Jan 04 2017;37(1):204-216. doi:10.1523/JNEUROSCI.2967-16.2016

2.	Kaiser EA, Kuburas A, Recober A, Russo AF. Modulation of CGRP-induced light aversion in wild-type mice by a 5-HT(1B/D) agonist. *The Journal of neuroscience : the official journal of the Society for Neuroscience*. Oct 31 2012;32(44):15439-49. doi:10.1523/JNEUROSCI.3265-12.2012
